# Comparative Transcriptomic Analysis of Obesity and Lipodystrophy Reveals Shared Mechanisms and Novel Targets in Adipose Tissue Dysfunction

**DOI:** 10.64898/2025.12.03.692243

**Authors:** Rola Shaaban, Antoine Rimbert, Yoann Combot, Simon Ducheix, Gilliane Chadeuf, Marie Palard, Mikael Croyal, Samy Hadjadj, Bertrand Cariou, Cedric Le May, Abdelhalim Larhlimi, Xavier Prieur

## Abstract

The global rise in obesity poses a major public health challenge. While chronic energy surplus is a well-established driver of weight gain and obesity, the mechanisms linking adipose tissue (AT) expansion to cardiometabolic complications remain incompletely understood. In obese individuals, dysfunctional AT loses its capacity to store excess lipids, leading to ectopic fat accumulation and contributing to cardiometabolic complications such as type 2 diabetes. However, the molecular events that drive the transition from healthy to dysfunctional adipocytes are poorly defined.

At the opposite end of the adiposity spectrum, lipodystrophies represent a heterogeneous group of disorders characterized by selective loss of AT, often accompanied by severe metabolic disturbances. Despite these contrasting adipose phenotypes, both obesity and lipodystrophy result in similar metabolic complications.

In this study, we investigated whether AT in these two contrasting conditions shares a common molecular signature. We performed an unbiased comparative transcriptomic analysis of AT from lipodystrophic BSCL2-deficient and obese mice, identifying a shared signature of 129 genes. Using publicly available datasets, we replicated this signature and refined it to 102 genes whose expression is consistently altered in both obese and lipodystrophic adipose tissue.

Correlation network analysis, gene ontology, and literature-based refinement revealed that these genes fall into nine functional categories: lipogenesis, adipocyte differentiation, carbohydrate metabolism, mitochondrial function, amino acid metabolism, reactive oxygen species, metabolic processes, immune response, and a group with no clear functional association. Most of these genes’ expression levels correlated strongly with insulin sensitivity across lipodystrophic and obese mice, as well as human samples.

Finally, 52 genetic loci containing these genes harbor variants associated with type 2 diabetes, including 11 loci where genetic associations directly influence candidate gene expression levels.

In conclusion, our findings demonstrate that a shared “energetic collapse” of adipocytes, characterized by profound metabolic inflexibility in pathways spanning glucose utilization, lipogenesis, and amino acid catabolism, represents a common pathogenic mechanism underlying adipose tissue dysfunction in both obesity and lipodystrophy. This convergent molecular signature underscores the critical role of intrinsic adipocyte metabolic health in systemic energy homeostasis and insulin sensitivity.

## Introduction

The global rise in obesity is a major public health concern and is closely associated with an increased risk of cardiometabolic diseases such as type 2 diabetes (T2D), metabolic dysfunction-associated steatotic liver disease (MASLD), kidney failure, and cardiovascular disease. While chronic energy surplus is a well-established driver of weight gain and obesity, the mechanisms linking adipose tissue (AT) expansion to cardiometabolic complications remain incompletely understood. Notably, not all individuals with obesity develop metabolic disease, as supported by epidemiological data and genome-wide association studies (GWAS), which have identified 62 *loci* that decouple body mass index (BMI) from metabolic risk^1^. Functional analyses suggest that these loci influence genes regulating adipocyte function and fat distribution. Moreover, *loci* associated with waist-to-hip ratio strongly correlate with insulin resistance, reinforcing the notion that impaired storage capacity in subcutaneous AT is detrimental to metabolic health^2^. Conversely, individuals with a genetic profile favoring subcutaneous fat accumulation often exhibit “favorable adiposity,” characterized by lower metabolic risk despite elevated fat mass^3^. Additionally, interindividual variability in weight gain, fat distribution, and metabolic outcomes in response to overfeeding suggests the existence of a personal fat threshold^4^, beyond which ectopic lipid deposition and metabolic disease develop^5^.

AT plays a central role in maintaining energy homeostasis. In lean individuals, radiotracer studies show efficient lipid storage post-prandially and release during fasting. In contrast, obese individuals display impaired lipid turnover, reflecting metabolic inflexibility^6^. At the cellular level, AT from obese patients exhibits altered glucose and lipid metabolism ^7^, alongside inflammation, fibrosis, hypoxia, and oxidative stress ^5^. Dysfunctional AT loses its ability to store excess lipids, promoting ectopic fat accumulation, particularly in the liver, which contributes to complications such as T2D and MASLD ^8^. While the consequences of AT failure are well documented, the underlying mechanisms remain elusive. In particular, the molecular events driving the transition from healthy to dysfunctional adipocytes are poorly defined. Gaining mechanistic insights into adipocyte dysfunction is essential for identifying novel therapeutic targets. This need is underscored by recent findings showing that obesity-related abnormalities persist in AT even after weight loss, due to epigenetic memory, emphasizing the importance of addressing intrinsic adipocyte dysfunction^9^.

At the other end of the adiposity spectrum, lipodystrophies are a heterogeneous group of disorders characterized by selective loss of AT, often accompanied by severe metabolic disturbances^10^. Congenital generalized lipodystrophy (CGL)—the most extreme form—is caused by mutations in four genes and is characterized by near-total absence of AT, leading to early-onset insulin resistance, MASLD, and cardiovascular disease ^11^. Preclinical studies have shown that the metabolic complications in mouse models of CGL stem from the absence of AT, and that restoration of adipose function—either by genetic rescue in adipocytes or transplantation of functional AT—can prevent insulin resistance and MASLD ^12^.

Despite their opposing adipose phenotypes, both obesity and lipodystrophy result in similar metabolic complications. Our group and others have studied mice lacking *Bscl2*, the gene responsible for the most severe form of CGL. These mice represent a unique model to investigate the metabolic consequences of extreme limitations in AT expandability, including lipotoxic and glucotoxic events in the liver, heart, and kidney ^13^. Importantly, while the metabolic complications overlap, the pathogenesis of AT failure differs between the two conditions: lipodystrophy reflects a primary defect in adipocyte biology, whereas in obesity, dysfunction arises secondarily to chronic overnutrition and adipocyte hypertrophy.

In this study, we sought to determine whether adipocyte dysfunction in these two contrasting conditions shares common molecular pathways. To this end, we performed an unbiased comparative transcriptomic analysis of AT from **BSCL2**-deficient and obese mice, followed by downstream analyses to identify shared pathways and novel candidate targets involved in AT failure.

## Results

### Transcriptomic analysis revealed a common signature in the T from lipodystrophic and obese mice

To test the hypothesis that different states of adipocyte dysfunction might share a common signature; we have compared AT from lipodystrophic mice and obese mice. For lipodystrophy, we used a model of inducible and progressive lipodystrophy; mice carrying a tamoxifen inducible deletion of *Bscl2*, the gene that encodes seipin, in the adipocytes, the iATSKO mice^14^. A 5-day tamoxifen treatment led to a decrease in inguinal fat pad weight from day 14 onward. As the subcutaneous adipose is the fat depot the most severely affected, this is the one that we studied to characterize the pathology. We have three groups of mice: 14 days, 1 month, 3 months after BSCL2 deletion. At 3 months, iATSKO mice presented with glucose intolerance and insulin resistance^14^. The AT transcriptomic profiles have been analyzed in an unbiased manner without prior biological assumptions, using the MaSigPro modeling tool to identify differentially expressed genes as a function of both time and phenotype. We identified 652 differentially expressed genes in the inguinal AT of iATSKO mice as compared to control littermates. These genes were grouped into nine clusters based on similar expression profiles (Supplemental Figure 1A). Three of these clusters exhibited particularly pronounced expression differences between individuals with iATSKO and controls. The first and second clusters revealed a progressive decrease in expression, from early to intermediate stages, of genes involved in mitochondrial function and lipogenesis, mainly (Figure 1A-1B and Supplemental Figure 2). The third cluster exhibited a median increase in expression, in iATSKO mice, of genes linked to endoplasmic reticulum stress (Figure 1C - Supplemental Figure 2).

**Figure 1.**
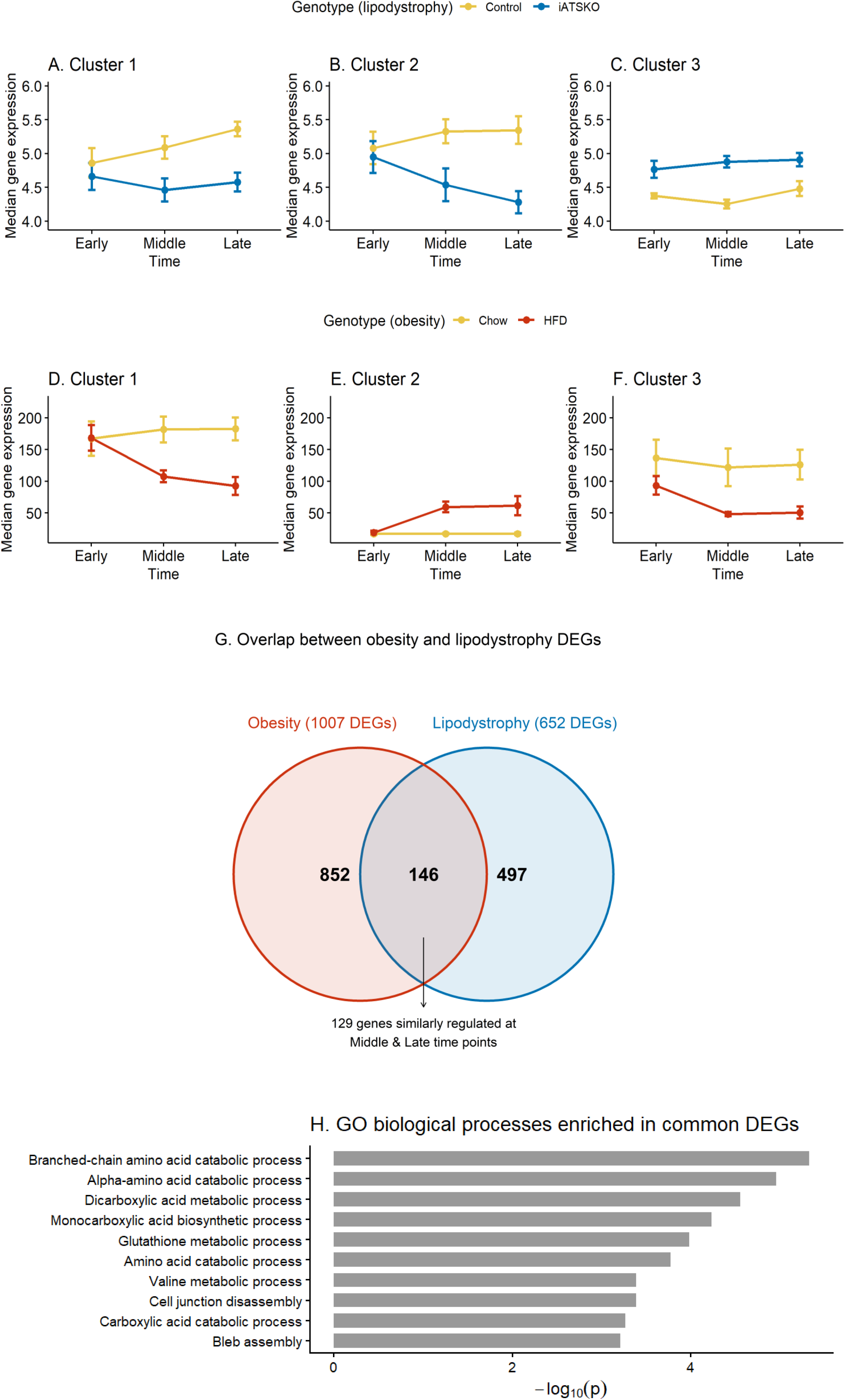
Identification of Differentially Expressed Genes in Adipose Tissue of Lipodystrophic and Obese Mice. (A–C) RNA sequencing analysis of subcutaneous adipose tissue from lipodystrophic iATSKO mice at 15 days (early), 30 days (middle), and 90 days (late) after tamoxifen injection. Gene expression data were clustered using MaSigPro, and the median expression profiles of three representative clusters (of nine total) are shown over time. (D–F) RNA sequencing analysis of visceral adipose tissue from C57Bl/6J mice fed either a chow diet or a high-fat diet (HFD) starting at 8 weeks of age, with samples collected at 1 month (early), 3 months (middle), and 6 months (late). MaSigPro clustering was applied, and the median expression profiles of three representative clusters (of nine total) are displayed over time. (G) Differentially expressed genes from both models were compared to identify commonly regulated genes. (H) Gene ontology analysis was performed on the list of commonly regulated genes.

For obesity, we used HFD diet feeding during 1, 3 or 6 months, and mice developed insulin resistance from 3 months. For the case of obesity, we used the gonadal AT, as this is the one that displays the most marked state of AT failure. MasigPro analysis identified 1007 differentially expressed genes as a function of phenotype and time. These genes were grouped into nine clusters according to their similar expression profiles (Supplemental Figure 1B). Two clusters (Figure 1D and 1F) exhibited a significant decrease in the median expression of genes involved in lipogenesis, as well as in various mitochondrial metabolic processes (Supplemental Figure 2). Conversely, the third cluster (Figure 1E) exhibited a median increase in the expression of genes associated with the immune system (Supplemental Figure 2).

We then performed a comparative analysis of differentially expressed genes in both models and identified 155 genes common to both datasets. We examined the regulation of these genes and retained only those that exhibited the same pattern—either overexpressed or downregulated—under both conditions at the median and late time points. This resulted in a list of 129 genes, which we consider a common gene signature between the two murine models of AT failure (Supplementary Table 1).

As an initial step to characterize their biological relevance, we conducted a Gene Ontology (GO) enrichment analysis. The results highlighted a strong enrichment for genes involved in mitochondrial function and energy metabolism, including pathways related to carbohydrate metabolism, lipogenesis and amino acid (aa) metabolism (Figure 1H). Altogether, this common transcriptional signature suggests a coordinated repression of key metabolic processes in dysfunctional AT, regardless of whether adipocyte failure arises from obesity or lipodystrophy.

### Gene Co-expression Network Analysis Reveals Conserved Metabolic Remodeling

To gain a deeper understanding of the altered metabolic pathways in adipocytes, we utilized the MEGENA (Multiscale Embedded Gene Co-expression Network Analysis) tool to construct gene-gene interaction networks and identify co-expression modules. Following network construction, Gene Ontology (GO) enrichment analysis was performed on each identified gene cluster to elucidate their predominant biological functions. These analyses were conducted in parallel on both the lipodystrophy and obesity datasets, revealing a remarkably conserved hierarchical organization of gene modules across both conditions. As depicted in Figure 1A (Lipodystrophy dataset), the 129 differentially expressed genes were initially aggregated into a Root module (M0) (Figure 2A). This root module subsequently resolved into two primary, well-separated branches Module 1 (M1), comprising 31 genes, was predominantly enriched for functions related to the immune system response (Figure Supplemental 3). The second major branch, Root module (M2), containing 84 genes, encompassed a broader range of adipocyte functions. Further hierarchical deconvolution of M2 revealed two distinct sub-branches: M3 with 44 genes, and M4 with 40 genes. From M3, two smaller modules were derived: M5, with 15 genes, was significantly enriched in carbohydrate metabolism pathways, while M6, consisting of 10 genes, was primarily associated with mitochondrial functions and amino acid catabolism (Figure Supplemental 3). From M4, M7, comprising 23 genes, was identified, and this module was specifically implicated in Reactive Oxygen Species (ROS) activity and various other metabolic processes. Additionally, 14 genes remained unclassified, not falling into any of the defined modules.

**Figure 2.**
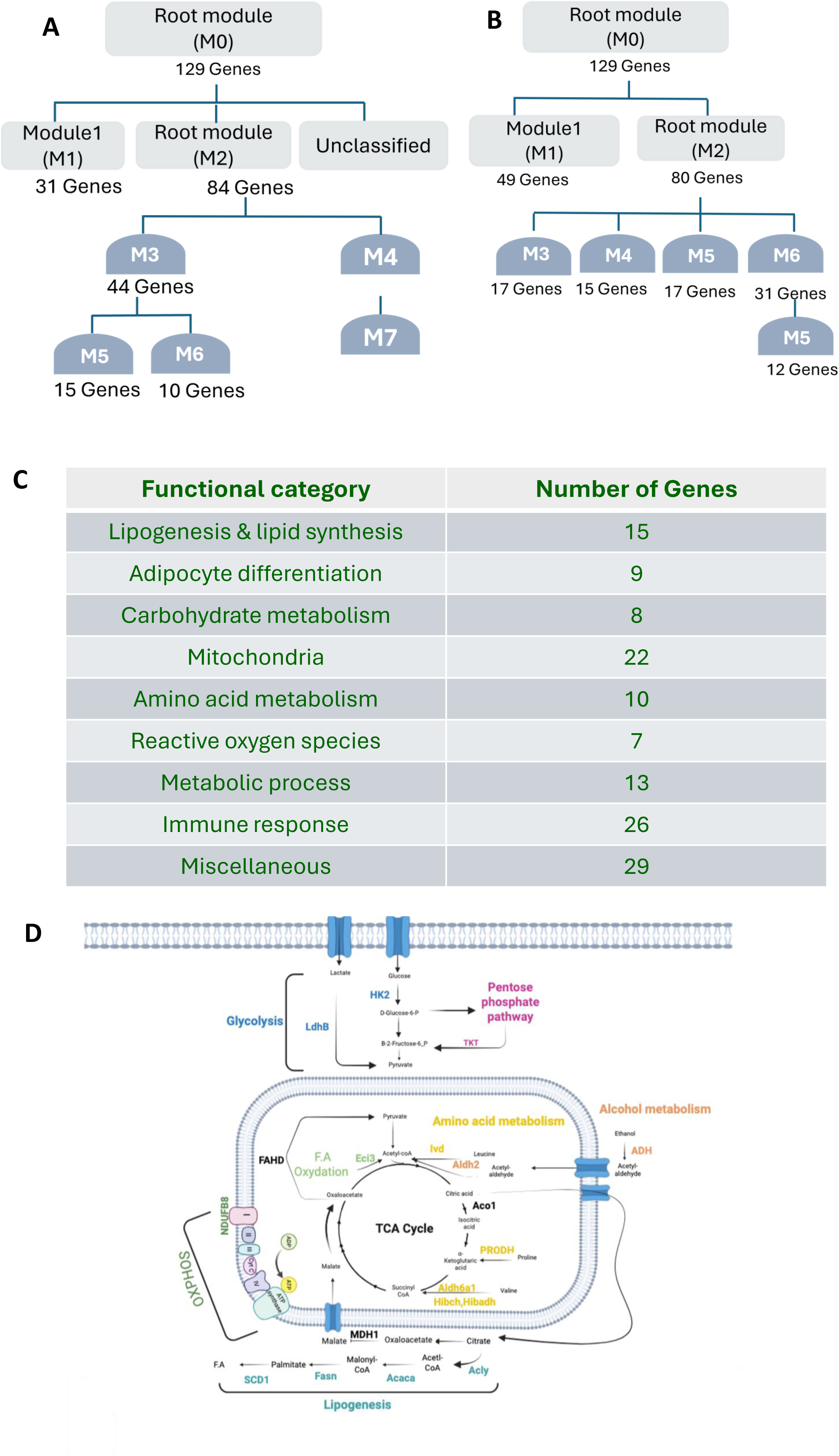
Multi-approach strategy to identify biological pathways in both AT failure conditions. MEGENA (Multiscale Embedded Gene Co-expression Network Analysisis used to construct gene-gene interaction networks and identify co-expression modules among the 129 genes in the lipodystrophy (A) and the obesity (B) dataset. Based on the GO analysis of each module and a comprehensive literature-based analysis, each genes has been classified into one functional category (C) and represented on a general bioenergetic cartoon of the adipocyte (D).

Similarly, in the obesity dataset, the same 129 genes were organized into a highly comparable hierarchical structure. Here, Module 1 (M1) contained 49 genes and, consistent with the lipodystrophy findings, was primarily involved in immune system response (Figure 2B). The Root module (M2) in obesity comprised 80 genes and further branched into four sub-modules. M3, with 17 genes, was associated with carbohydrate metabolism. M4, also with 17 genes, was enriched in mitochondrial functions and amino acid catabolism. M5, comprising 15 genes, was linked to ROS activity and various metabolic processes. Finally, M6, containing 31 genes, and its derived sub-module M7, with 12 genes, were both strongly associated with fat cell differentiation. The observed preservation in the overall architecture and functional enrichments of the gene modules across both lipodystrophy and obesity datasets strongly suggests shared underlying mechanisms of adipocyte dysfunction.

Based on the GO analysis of each module and a comprehensive literature-based analysis, we classified the identified genes into nine functional categories, as summarized in Figure 2C. Specifically, the major functional categories identified were: Lipogenesis & lipid synthesis, Adipocyte differentiation, Carbohydrate metabolism, Mitochondria, amino acid metabolism, Reactive oxygen species (ROS), Metabolic process and the immune response. Additionally, we identified a group of genes without a clear common biological function, which we categorized as “miscellaneous.”

To illustrate the interplay between the affected metabolic pathways, we generated a schematic representation integrating the 19 metabolism-related genes identified (Figure 2D). At a glance, the common metabolic signature reveals that in dysfunctional adipocytes, the expression of genes involved in glucose and amino acid catabolism is decreased. These two pathways supply the TCA cycle, and the expression of genes related to this cycle are also repressed. Finally, the mRNA levels of genes responsible for fatty acid synthesis from citrate are likewise reduced.

### Toward a robust signature of adipocyte dysfunction

In order to strengthen our findings, we tested our 129 signature-genes in 4 transcriptomic publicly available datasets of visceral adipose tissue of mice fed a HFD (detailed in methods section and in Figure 3). We analysed each data set to search for differentially expressed genes with the R package LIMA for the microarray data set (GSE97271) and with R package **edgeR** for RNA sequencing data (GSE213632, E-MTAB-8840 and GSE167264).

**Figure 3.**
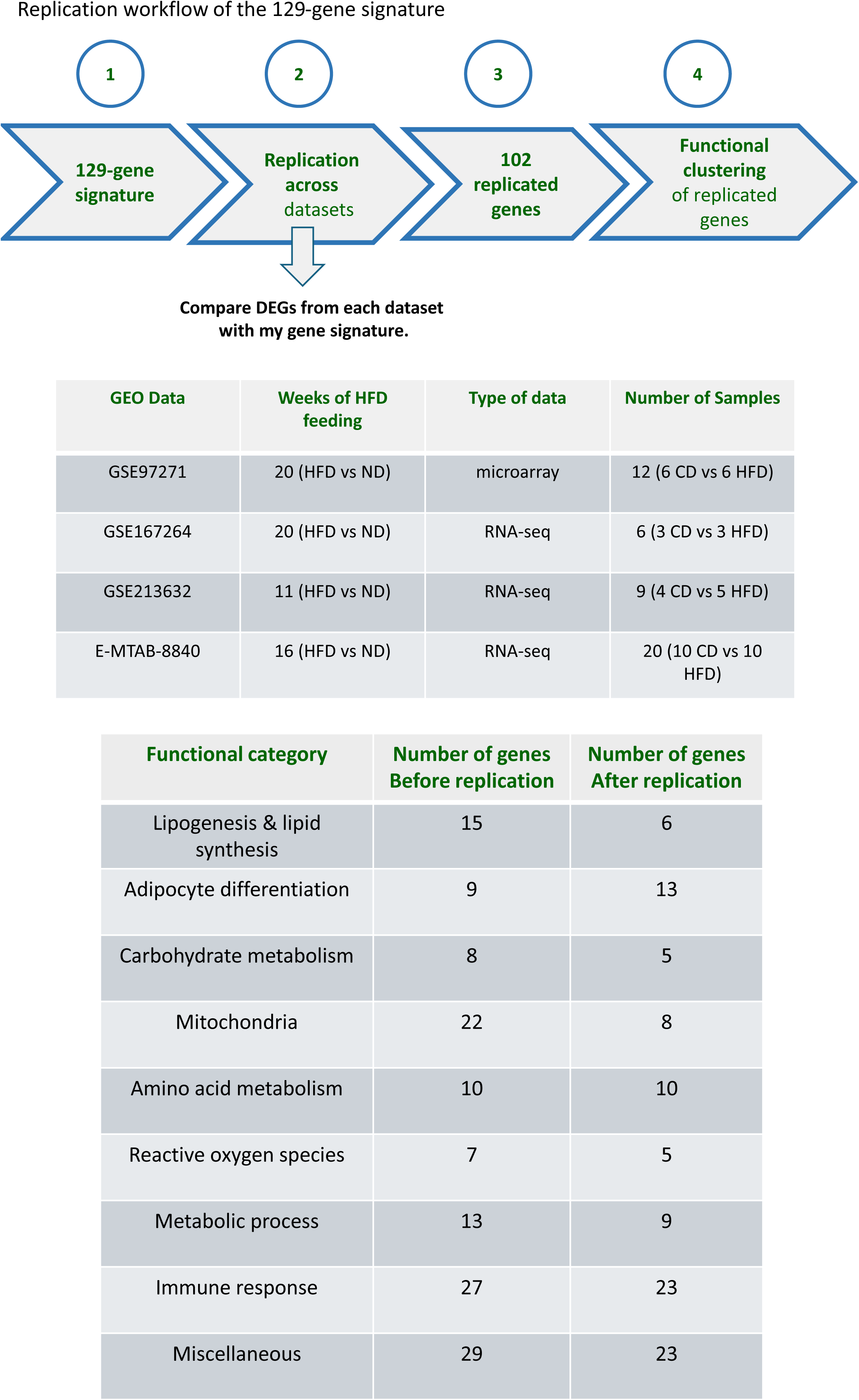
Identification of genes consistently altered in adipose tissue (AT) dysfunction across multiple datasets. Differential gene expression analysis was performed using four publicly available transcriptomic datasets of visceral adipose tissue from high-fat diet (HFD)-fed mice. In each dataset, genes differentially expressed between lean and obese conditions were identified and compared to the shared gene signature highlighted in Figure 1. The orange table indicates 102 genes that are consistently dysregulated across these independent AT dysfunction datasets.

Then, we tested the behavior of our 129 genes and selected the genes whose expression was similarly affected in at least 3 of the data sets. We retrieved 102 of these genes that were repeatedly differentially expressed in obese mice compared to lean controls. This replication reinforces the importance of these genes individually. Table 1 lists the 102 genes constituting the adipocyte dysfunction signature, along with their corresponding functional categories. Figure 3 provides a summary of the number of genes associated with each biological category. From this point onward, all analyses will be based on this 102-gene signature.

**Table 1:**
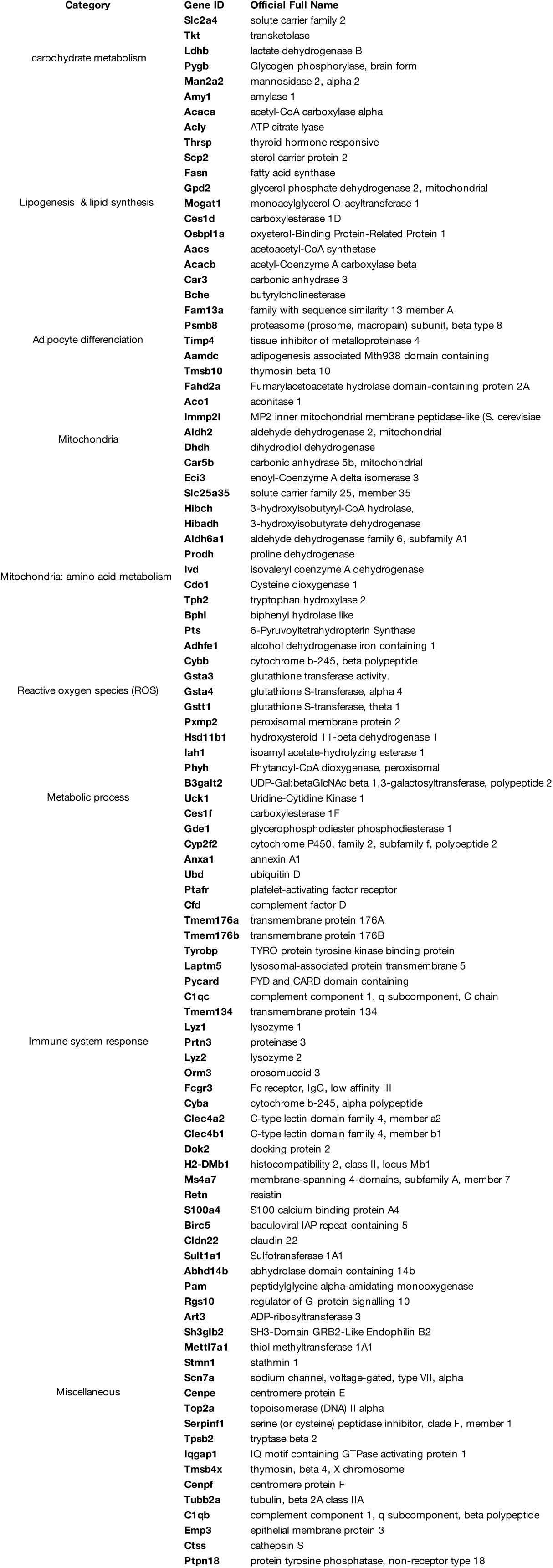
Adipocyte dysfunction gene signature. This table presents the 102 genes repeatedly identified as differentially expressed in obese mice compared with lean controls, defining a robust adipocyte dysfunction signature. The recurrence of these genes across datasets reinforces their individual relevance to obesity-associated alterations in adipocyte biology. Each gene is listed together with its assigned functional category, highlighting the major biological processes perturbed in dysfunctional adipocytes.

### The expression levels of genes involved in amino acid catabolism and lipogenesis are strongly correlated to insulin sensitivity, in mice and human

To improve the biological interpretation of our data, we performed weighted correlation network analysis (WGCNA) between geane expression and insulin resistance, represented here by insulin tolerance test and glucose tolerance test (GTT) values. This method allows the identification of co-expressed gene clusters and their correlation with a phenotypical trait. For the lipodystrophy data set, we identify a cluster of 76 genes that were negatively correlated with the GTT AUC values (Figure 4 A-B). This means that low expression of these genes is associated with high AUC values and therefore poor glucose tolerance. Running GO analysis on these genes, the top terms associated with these genes were all linked to lipogenesis and amino acid metabolism (Figure 4C). On the other hand, a second cluster gathers 16 genes that were not co-expressed but whose expression was positively correlated with the GTT AUC values, meaning that their high expression was positively associated with poor glucose tolerance. Interestingly, upon the 102 gene of the common signature, 92 were correlated, positively or negatively, with the AUC values.

**Figure 4.**
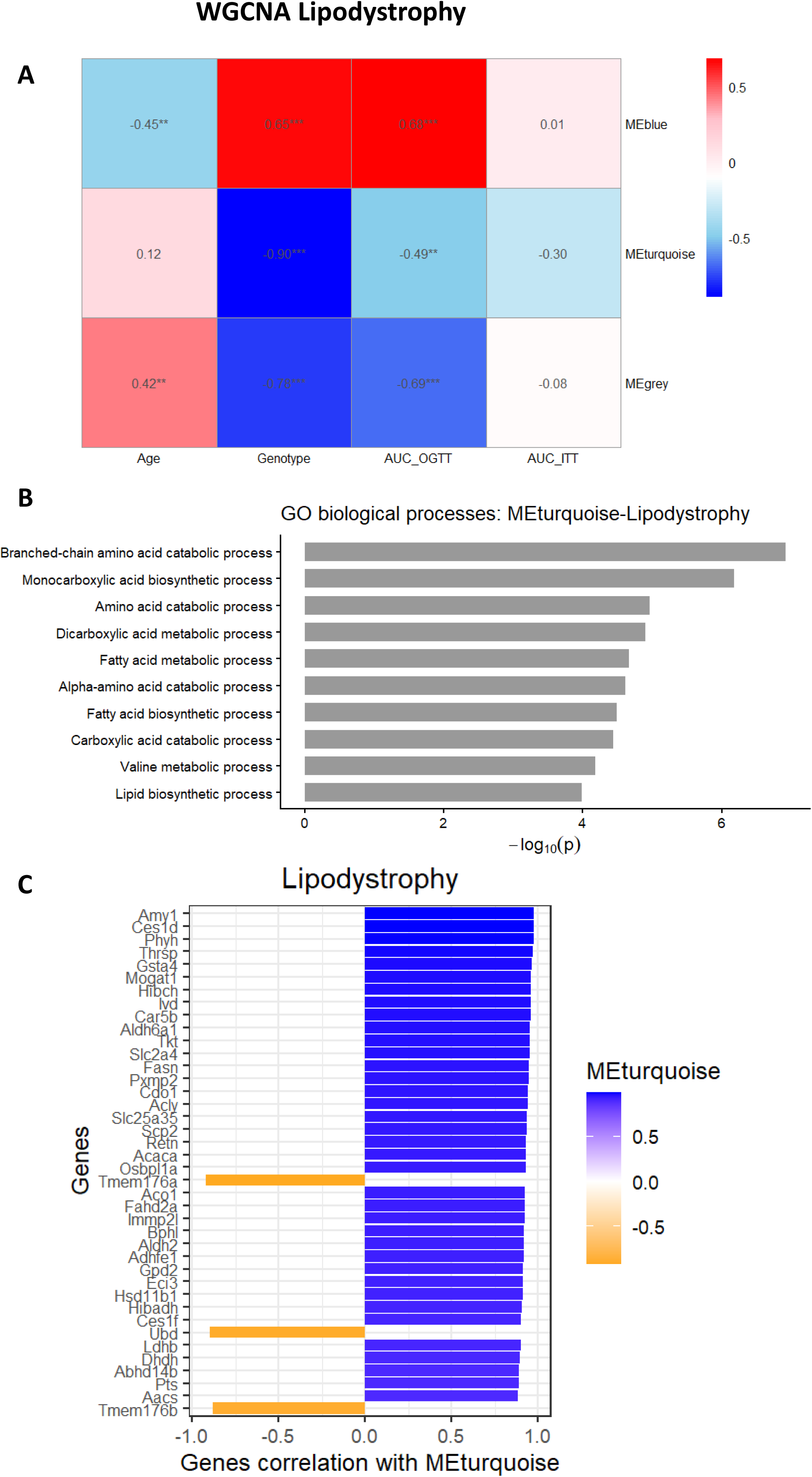
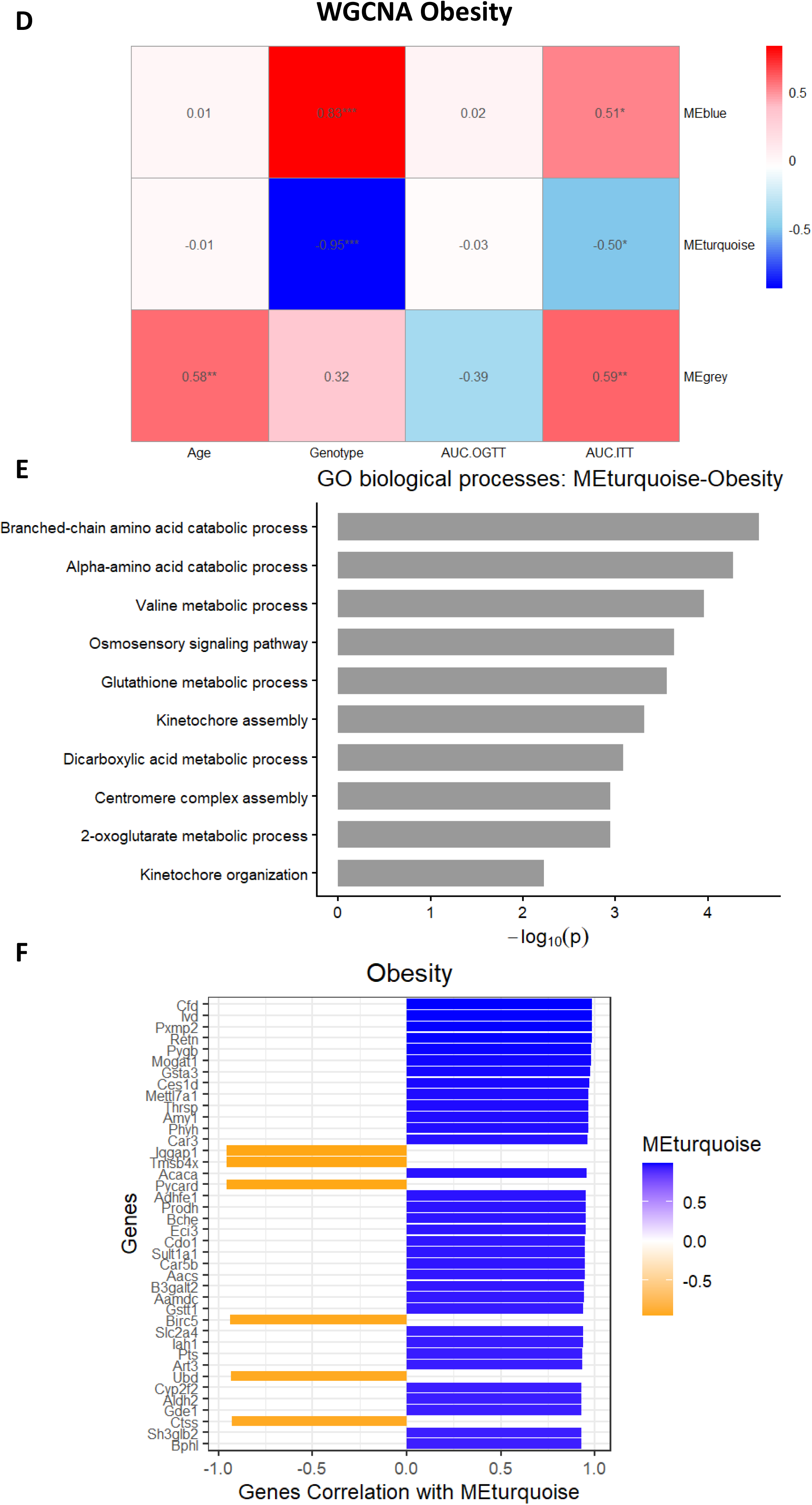
Weighted Gene Correlation Network Analysis (WGCNA) of Gene Expression and Metabolic Status. WGCNA was performed on the list of replicated genes using data from lipodystrophic (A–C) and obese (D–F) mouse models. Correlations between gene expression and metabolic traits, including insulin sensitivity (AUC ITT) and glucose tolerance (A, D), were assessed. In each model, the association between the turquoise module and metabolic traits was evaluated (B, E). Gene ontology analysis was conducted to identify functional categories most strongly correlated with insulin sensitivity (C, F)

For the obesity dataset, the profile was similar, with a turquoise cluster gathering 83 genes inversely correlated with GTT AUC values and only 8 genes in the grey cluster positively associated with glucose intolerance (Figure 4D-E). Using GO analysis with the turquoise gene, again, the major pathways highlighted were the lipid synthesis pathway, including lipogenesis, the amino acid catabolism pathway, including the Branched-chain amino-acids (BCAA) metabolism, and the TCA cycle (Figure 4F). Altogether, in two mouse models of AT failure, and in 2 different AT fat pads, glucose intolerance is associated with decreased expression of genes involved in TCA cycle, lipogenesis, and aa metabolism.

To assess the relevance of our shared gene signature in humans, we investigated whether the expression levels of our 102-gene list were correlated with key metabolic traits, including body mass index (BMI), waist-to-hip ratio, HOMA-IR, HbA1c, glycemia, and insulin levels.

For this analysis, we utilized the recently released adipose tissue portal, which compiles a comprehensive dataset enabling such correlation studies^15^. We assessed the correlation of our 102-gene list with metabolic traits using transcriptomic data from subcutaneous adipose tissue (Figure 5 and Table 2), and performed the same analysis in visceral adipose tissue (Supplemental Figure 4). The correlations between the expression levels of our 102-gene set and the phenotypic traits are more pronounced in subcutaneous adipose tissue, which includes a larger number of subjects (on average ∼50 individuals) compared with visceral adipose tissue (approximately ∼10 individuals). Genes were classified into functional categories as previously defined. In subcutaneous adipose tissue, the expression of 76 genes was significantly correlated with at least four of the phenotypic traits examined (Table 2). The functional classification of these 76 genes is presented in Table 3, highlighting the main biological categories associated with their expression profiles.

**Figure 5.**
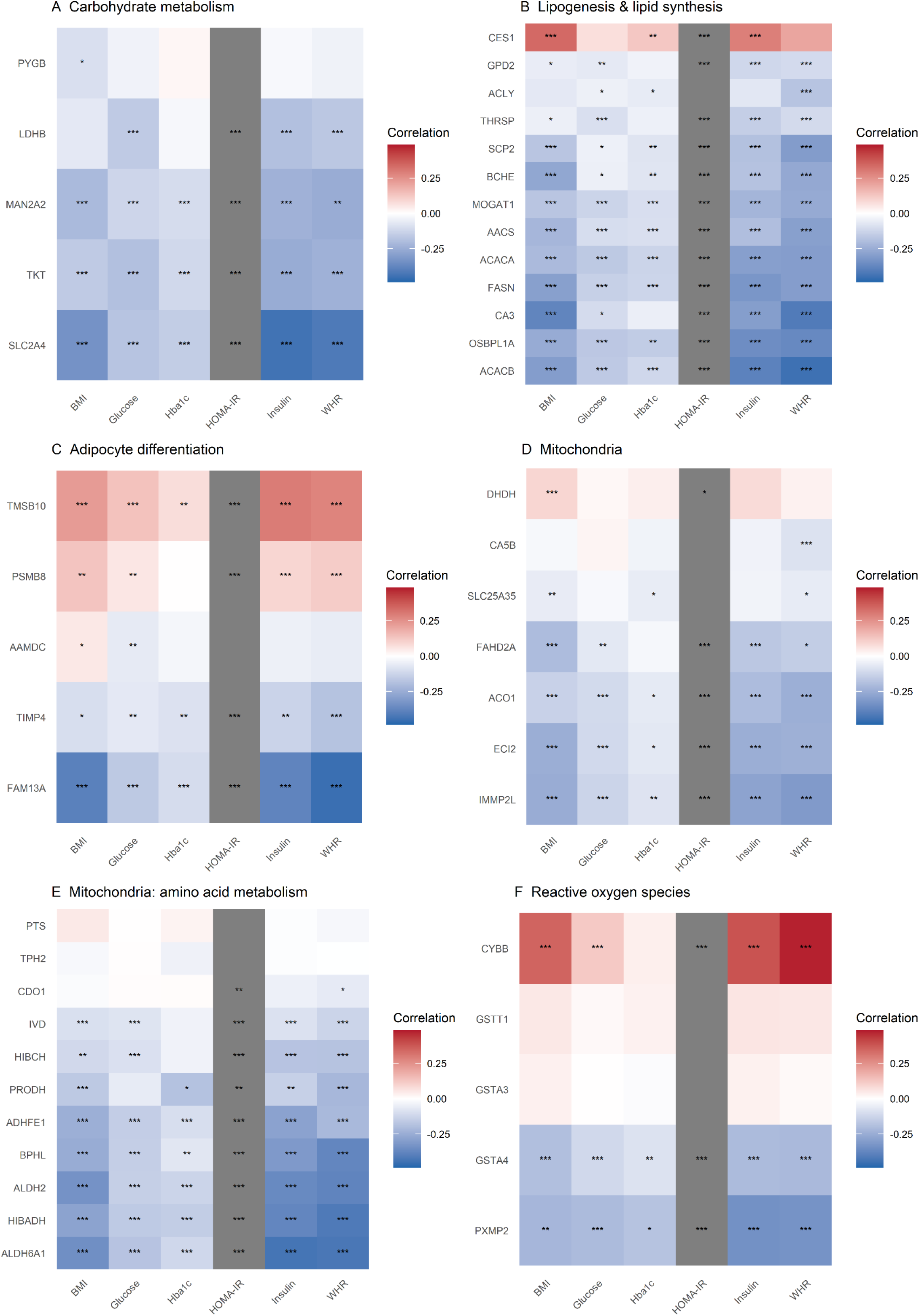
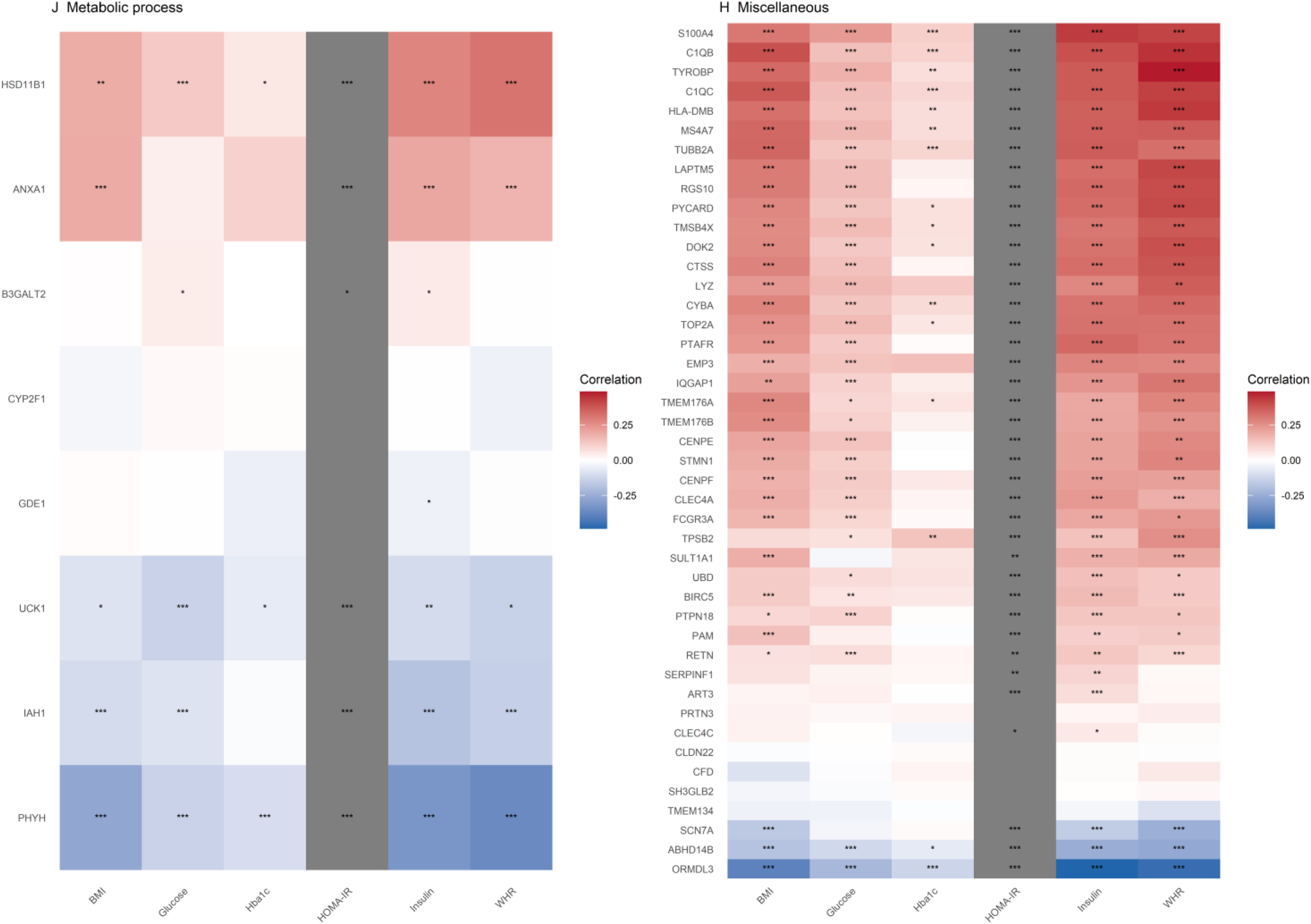
Correlation of mRNA Levels of Commonly Altered Genes with Phenotypic Traits in Humans. The replicated list of commonly altered genes was analyzed using the publicly available Adipose Tissue Portal to assess correlations between gene expression and phenotypic traits, including BMI, HbA1c, HOMA-IR, plasma insulin levels, and hip-to-waist ratio (HWR). Each heat map displays genes from a specific functional category, as defined in Figure 2. Statistical significance was determined using the Mann-Whitney test (*p < 0.05, ***p < 0.001, ****p < 0.0001).

**Table 2:**
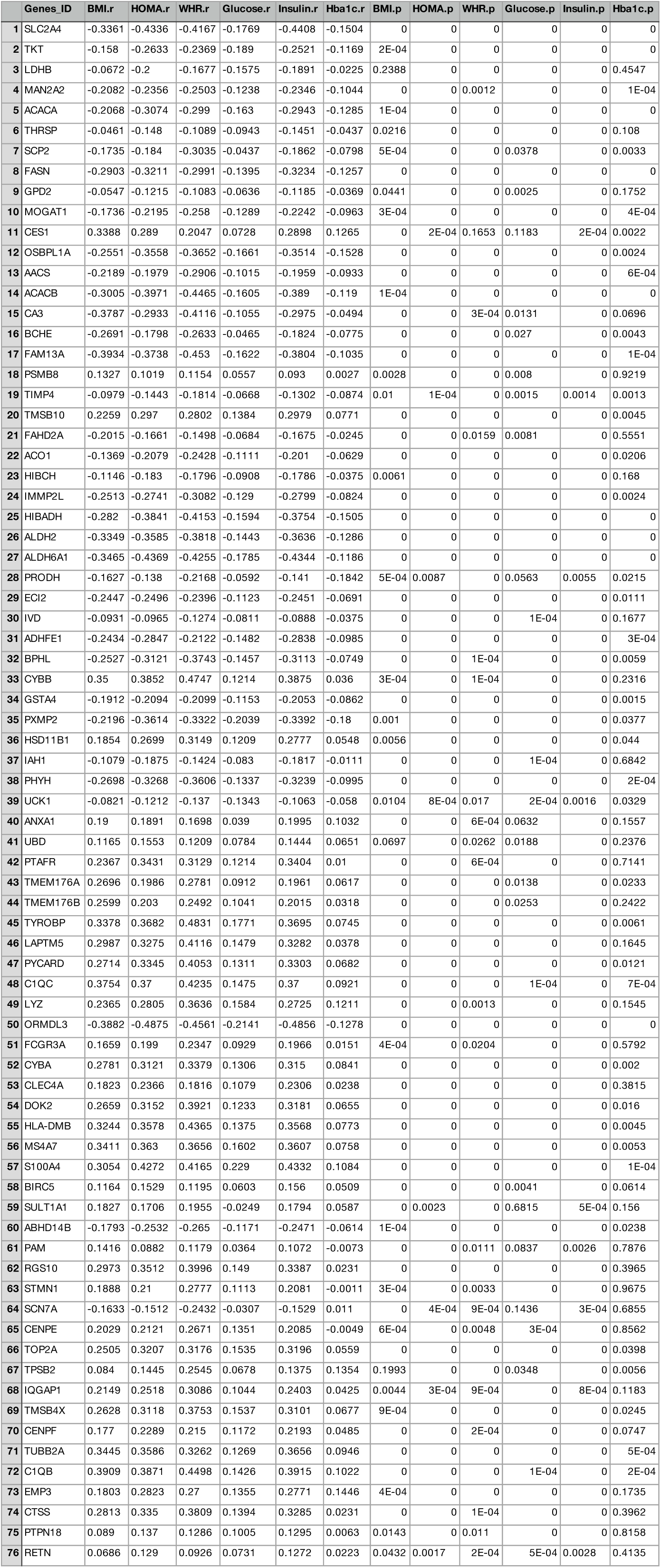
Correlation between the mRNA levels of the indicated genes in subcutaneous adipose tissue from iATSKO mice and the phenotypic traits listed, calculated using the WGCNA pipeline.

**Table 3:**
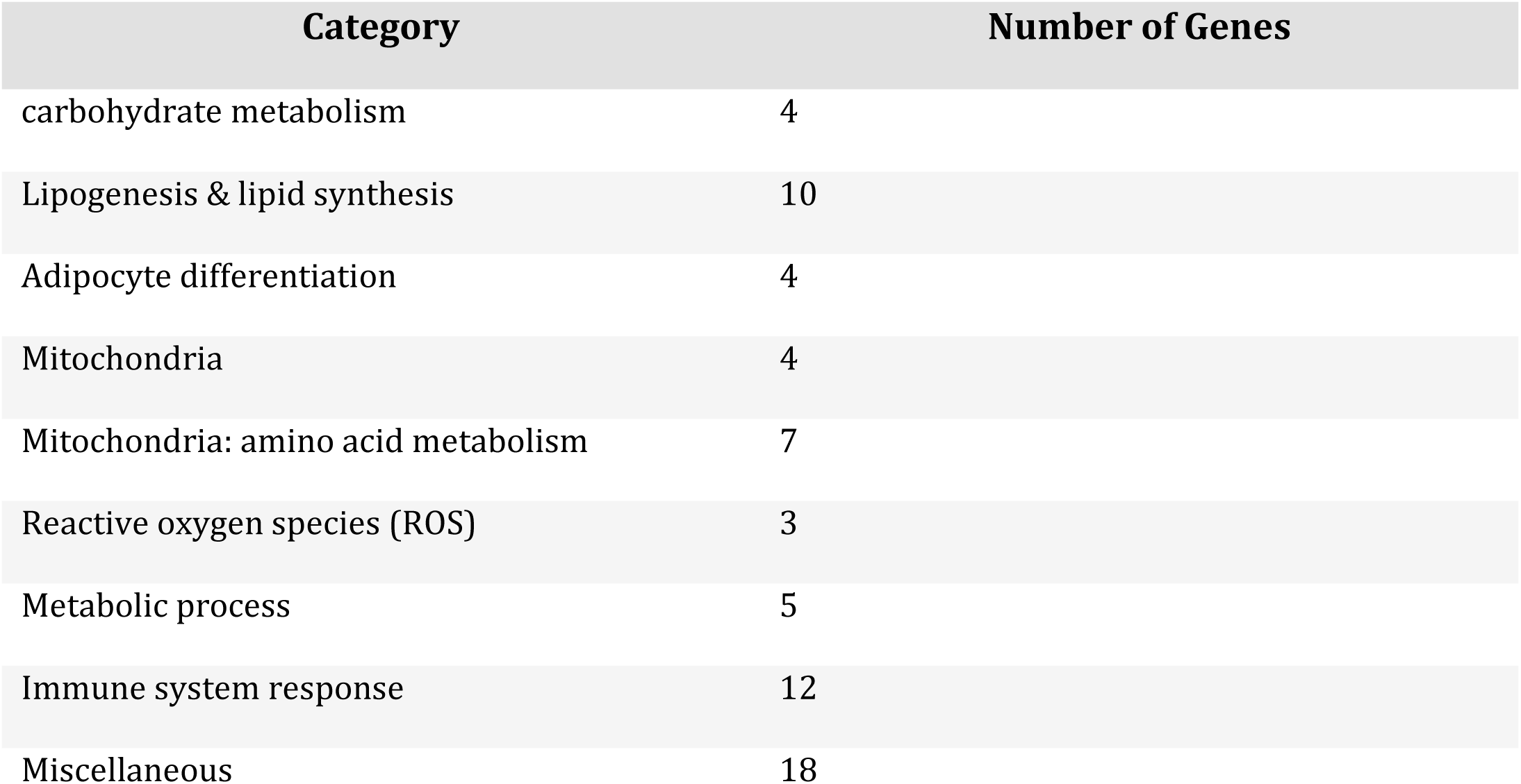
Functional classification of the 76 genes whose expression in subcutaneous adipose tissue was significantly correlated with at least four phenotypic traits (as identified in Table 2). Genes are grouped according to major biological processes and functional categories associated with their expression profiles.

| Category | Number of Genes |
| --- | --- |
| carbohydrate metabolism | 4 |
| Lipogenesis & lipid synthesis | 10 |
| Adipocyte differentiation | 4 |
| Mitochondria | 4 |
| Mitochondria: amino acid metabolism | 7 |
| Reactive oxygen species (ROS) | 3 |
| Metabolic process | 5 |
| Immune system response | 12 |
| Miscellaneous | 18 |

### Genome-wide association analysis highlights potential contributors to adipocyte dysfunction

Then we ask whether the function of these genes could be directly related to the common phenotypical trait between lipodystrophy and obesity, which the development of T2D. We screened for evidence associating common human genetic variants in the *loci* of our 102 candidate genes with T2D in large population-based cohort studies ^16^. Among our 102 gene-*loci* 52 presented with significant genetic association (p<5E-8) with T2D (Table 4). In order to refine the associations in genetic *loci* of interest, we tested the association of identified variants with the expression of nearby genes (including candidate genes) in AT, using expression quantitative trait *loci* datasets ^17,18^. The locus-refinement led to the identification of variants significantly associated with: T2D on the one hand, and with changes in gene-expression of 11 candidate genes in AT on the other hand, supporting the causal association of the gene with T2D in the *locus* (Table 5).

**Table 4:**
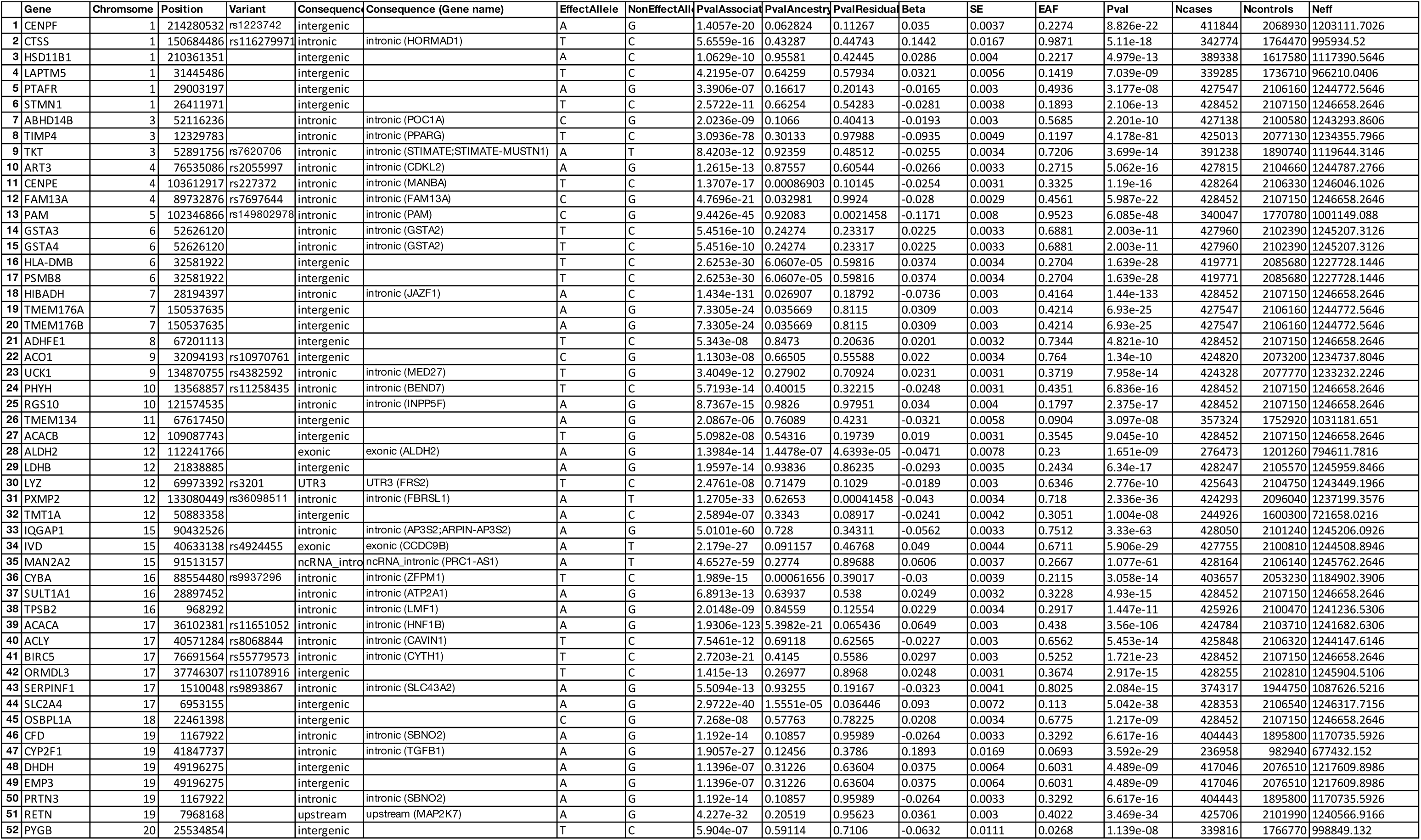
Genetic association analysis of the 102 gene loci. Fifty-two loci showed a significant association with type 2 diabetes (p < 5×10⁻⁸).

**Table 5.** Locus-refinement analysis integrating genetic association and adipose tissue eQTL data. Variants significantly associated with T2D and with the expression of 11 candidate genes in adipose tissue are shown, supporting their putative causal role within the corresponding loci.

| Gene | SNP | P value SAT<br>ref 18 | P value GTEX<br>SAT | P value GTEX<br>VAT |
| --- | --- | --- | --- | --- |
| PXMP2 | 12_133080449_A_T | 5.724217e-09 | 0.064 | 0.97 |
| LDHB | 12_21838885_G_A | 0.0025591 | 0.28 | 0.41 |
| PHYH | 10_13568857_T_C | 0.0300354 | 0.31 | 0.88 |
| FAM13A | 4_89732876_C_G | 5.790465e-66 | 3.3e-12 | 0.011 |
| Sult1A1 | 16_28897452_G_A | 1.33996e-05 | 2.2e-8 | 5.8e-8 |
| Adhfe1 | 8_67201113_T_C | 9.374944e-41 | N/A | N/A |
| Ivd | 15_40633138_A_T | 2.384192e-07 | 0.58 | 0.79 |
| MAN2A2 | 15_91513157_T_A | 0.0001317322 | 0.015 | 0.14 |
| PYGB | 20_25534854_C_T | 0.03543784 | 0.20 | 0.043 |
| ORMDL3 | 17_37746307_T_C | 1.495861e-25 | 0.059 | 0.032 |
| TIMP4 | 3_12329783_C_T | 2.644614e-33 | N/A | N/A |

Finally, Table 6 presents a bibliographical analysis of the biological functions of genes and their involvement in the pathogenesis of obesity and its associated complications. Among the prioritized genes, we have identified *FAM13A*^1^ and *SULT1A1*^19^ which have already been implicated in obesity. Beyond the identification of known genes involved in AT dysfunction, which validates our findings, our study suggests the involvement of additional ones with unknown functions (e.g. *ORMDL3, ADHFE1; PXMP2; TIMP4, IVD, CENPE, PYGB, PHYH*, and *LDHB*. Further research is warranted to functionally test their involvement in AT dysfunction.

**Table 6.** Bibliographical review of the biological functions of prioritized genes and their involvement in obesity and related metabolic complications. Known genes previously implicated in adipose tissue dysfunction (e.g., *FAM13A*, *SULT1A1*) are highlighted, along with additional candidates of unknown function whose potential roles warrant further investigation.

| Gene Name | Symbol | Obesity/Metabolism Relevance |
| --- | --- | --- |
| Isovaleryl-CoA dehydrogenase | IVD | Decreased expression in obesity; SNPs associated with T2D and gene expression. Leucine catabolism and IVD activity are highly induced during adipogenesis, but its role in adipocytes remains poorly studied.<sup>48</sup> |
| Alcohol dehydrogenase, iron containing 1 | ADHFE1 | Poorly studied in adipose tissue; mRNA induced during adipocyte differentiation. Overexpressed in cancer, conferring metabolic flexibility; its role in adipocytes is speculative.<sup>51,52</sup> |
| Family with sequence similarity 13, member A | FAM13A | Modulates insulin sensitivity and may worsen high-fat diet effects; not essential for normal adipocyte insulin response, but overexpression promotes apoptosis. GWAS links FAM13A to favorable adiposity (more fat, lower cardiovascular risk).<sup>1,25,26</sup> |
| Tissue inhibitor of metalloproteinase-4 | TIMP4 | Facilitates adipocyte differentiation in vitro; knockout protects mice from diet-induced obesity, possibly via reduced intestinal lipid absorption<sup>28</sup>. |
| Peroxisomal membrane protein 2 | PXMP2 | Highly expressed in mammary fat pad; deletion leads to abnormal lipid metabolism. Its specific role in adipocytes is not yet defined<sup>29</sup>. |
| Sulfotransferase family 1A | Sult1a | Deletion in adipocytes protects mice from diet-induced obesity, suggesting a protective role in adipocyte homeostasis<sup>19</sup>. |
| Phytanoyl-CoA hydroxylase | Phyh | Involved in glycogen metabolism, but not studied in obesity or adipose tissue. |
| Mannosidase alpha class 2A member 2 | Man2A2 | Not studied in adipocytes, but GWAS links SNPs to fasting insulin levels, suggesting a possible metabolic role<sup>56</sup>. |
| Glycogen phosphorylase, brain form | PYGB | Largely uncharacterized; GWAS associates variants with childhood obesity, indicating a potential role in early metabolic regulation<sup>57</sup>. |
| Lactate dehydrogenase B | LDHB | Key enzyme in glucose catabolism; little is known about its function in adipose tissue. |
| ORM1-like protein 3 | ORMDL3 | Primarily involved in immune system regulation; limited relevance to adipose tissue biology. |

## Discussion

Nearly two decades ago, AT expandability hypothesis proposed that the capacity to store excess calories as lipid in AT is limited and determined, for each individual, by the interplay of genetic and environmental factors ^20^. When this expansion limit is reached—most commonly in individuals living with overweight or obesity—excess fat is redirected to non-adipose organs such as the liver^21^. This ectopic fat deposition promotes lipotoxicity and contributes to the development of MASLD and T2D. The hypothesis is supported by evidence from both human and animal studies: individuals with obesity who preferentially store fat in subcutaneous AT^22^, as well as obese mice with increased subcutaneous AT expansion^23^, are relatively protected from the development of T2D and MASLD. Conversely, patients with lipodystrophy ^10^ and mouse models characterized by limited adipose storage capacity develop severe metabolic complications^12,24^. The lipodystrophy paradigm has therefore been widely used to investigate the mechanisms underlying lipotoxicity in non-adipose organs^11^. Despite these advances, the origins of adipocyte dysfunction remain poorly understood. We sought to determine whether studying lipodystrophy could provide insights into why and how adipocytes become dysfunctional. To our knowledge, this is the first study to directly compare the transcriptomic landscape of AT failure in the context of both obesity and lipodystrophy. We identified 132 genes that are commonly dysregulated in both conditions. Using a replication strategy, we validated the dysregulation of 102 of these genes in four independent datasets of AT isolated from obese mice. Furthermore, analysis of human data confirmed that a subset of these genes is correlated with metabolic complications. Notably, 52 genetic *loci* containing our genes of interest harbor variants associated with T2D. Among them we identified 11 genetic association directly linked to candidate-genes as they influence their gene expression levels. Collectively, these findings provide new insights into the molecular basis of AT dysfunction and its role in the pathogenesis of metabolic diseases.

Among the 11 genes prioritized by our genetic studies, FAM13A stands out as a gene of considerable interest, having already attracted attention in the context of obesity. Experimental studies in animal models suggest that FAM13A influences insulin sensitivity and may exacerbate the metabolic consequences of a high-fat diet^25^. However, the picture is nuanced: while one study found that FAM13A is not essential for the normal insulin response in adipocytes, overexpression of this gene in these cells appears to promote apoptosis^26^. Adding to this complexity, a recent genome-wide association study identified *FAM13A* as a potential promoter of favorable adiposity—characterized by increased fat mass but a reduced risk of cardiovascular disease^1^. *TIMP4, PXMP2* and *SULT1A* seem to be involved in adipocyte differentiation or homeostasis, but further studies are necessary to properly underatnd how they control adipocyte phenotype. *TIMP4* (Tissue Inhibitor of Metalloproteinase-4) has been shown to facilitate adipocyte differentiation *in vitro*^27^. In mice, global knockout of *Timp4* confers protection against diet-induced obesity, an effect that appears to be mediated by decreased intestinal lipid absorption rather than direct action within adipocytes^28^. PXMP2 is another gene of interest, given its high expression in the mammary fat pad and the observation that its deletion leads to abnormal lipid metabolism^29^. Recent research indicates that deletion of *Sult1a* in mouse adipocytes can protect against the deleterious effects of diet-induced obesity, pointing to a possible protective function that warrants deeper investigation^19^. These findings validate our approach in identifying promising targets that play a central role in adipose tissue homeostasis. Continued functional exploration of these genes is warranted to further elucidate their importance and potential as therapeutic targets.

Then, we sought to interpret our results by examining the modulation of pathways and biological processes. Our data were generated from bulk RNA sequencing, which, while adding complexity to the interpretation—since observed changes may originate from various AT cell types—also offers significant advantages. Notably, bulk RNA sequencing minimizes tissue manipulation that could alter the transcriptome, such as collagenase dissociation, and provides deep and robust sequencing coverage. Importantly, several single-nucleus RNA-seq studies in obese humans and mice have confirmed that genes involved in adipocyte metabolic function (including carbohydrate metabolism, amino acid catabolism, mitochondrial activity, and lipogenesis)—which we found to be downregulated—are specifically repressed within adipocytes themselves, thereby reinforcing the cell-autonomous interpretation of our findings ^30^. By integrating clustering and annotation strategies, we identified 9 pathways commonly altered in both models. Among these, immune system activation was evident, with a markedly higher number of genes involved in the context of obesity, consistent with extensive literature underscoring the role of inflammation in AT maladaptation during chronic overnutrition ^31^. However, previous studies have demonstrated that the immune response in the AT of lipodystrophic models differs from that observed in obese mouse AT^32^. Consequently, we did not focus our analysis on immune remodeling. Additionally, although fibrosis and hypoxia have been extensively investigated as drivers of AT failure ^33,34^, neither was identified as commonly altered in both of our models. Thus, given our comparative paradigm of two AT failure models, our work specifically emphasizes the metabolic remodeling occurring within adipocytes. This includes changes in carbohydrate metabolism, amino acid catabolism, mitochondrial function, diverse metabolic processes, lipogenesis, and ROS production. Importantly, our WGCNA revealed that alterations in these pathways are positively correlated with the development of insulin resistance, a finding further corroborated by analyses using the AT Portal in human datasets.

Previous studies have shown that genes related to carbohydrate metabolism, the TCA cycle, amino acid catabolism, and lipogenesis are strongly downregulated in the AT of women with metabolic syndrome or obesity compared to lean controls ^35^. Notably, bariatric surgery and the resulting weight loss induce a re-expression of genes associated with these same pathways ^36^. Multi-omic analyses, integrating lipidomics and transcriptomics, have demonstrated that in the AT of obese individuals, adipocyte hypertrophy is inversely correlated with the expression of TCA cycle and lipogenic genes and their products ^37^. Furthermore, genome-scale metabolic models (GEMs) based on transcriptomic data have highlighted that the TCA cycle, amino acid metabolism—particularly BCAA and tryptophan—and lipogenesis are strongly repressed in the AT of obese individuals ^38,39^. Recent advances in single-nucleus analysis of AT have identified distinct adipocyte subtypes and revealed that obesity drives remodeling of these subtypes, promoting adipocytes with cellular stress features and reducing the proportion of highly lipogenic adipocytes ^40^.

Our data demonstrate for the first time that this profound downregulation of key metabolic pathways in adipocytes is a shared feature of both hypertrophic adipocytes resulting from positive energy balance, as seen in obesity, and primary adipocyte dysfunction, such as lipodystrophy due to seipin deficiency. Strikingly, in both scenarios, adipocytes appear to enter a metabolically “frozen” state, characterized by a marked slowdown in pathways ranging from glucose catabolism and citrate production to fatty acid synthesis, which is the most efficient mechanism for lipid storage. The strong correlation between suppressed lipogenesis and insulin resistance raises important questions, as our data suggest that the loss of lipogenic capacities in adipocytes may underlie systemic metabolic dysfunction. This is consistent with mechanistic studies showing that inhibition of glucose uptake in adipocytes impairs lipogenesis and adversely affects systemic insulin sensitivity ^41^. Indeed, the glucose-sensitive transcription factor ChREBP serves as a key integrator of glucose utilization, lipogenesis, and insulin sensitivity in adipocytes ^42^. While the association between impaired lipogenesis and insulin resistance is strongly supported by previous studies ^43^and reinforced by our own results, it remains correlative. Targeted functional manipulation of the affected pathways will be required to establish causality and delineate the precise contribution of metabolic inflexibility to systemic insulin resistance. Therefore, our data suggest that supporting the entire metabolic pathway from glucose utilization to lipogenesis may help prevent adipocyte dysfunction in both primary and secondary forms of adipose tissue failure.

In addition to lipogenesis, aa metabolism consistently emerged as a significant functional category across all our analyses. Notably, the expression levels of several genes involved in AA metabolism were strongly correlated with insulin resistance, both in our murine models and in the adipose tissue of human patients living with obesity. The link between AA metabolism and obesity has long been recognized, dating back to early observations that BCAA leucine, isoleucine, and valine—are elevated in the plasma of obese individuals and correlate with insulin levels, suggesting a connection to insulin resistance^44^. Some studies have further shown that BCAA supplementation alone can induce insulin resistance, primarily through activation of the mTOR and JNK signaling pathways in skeletal muscle^45^. In adipose tissue, obesity is typically associated with reduced BCAA catabolism, and this pathway has been implicated in the regulation of adipogenesis^46^. While BCAAs are known to activate mTORC1 signaling, the precise contribution of mTORC1 to BCAA-mediated effects on adipocyte homeostasis in obesity remains to be fully clarified. Within our gene signature, we observed decreased expression of Ibadh and Hibch, two enzymes involved in valine catabolism. Previous research has shown that knockdown of these enzymes impairs glucose uptake and insulin signaling in adipocytes, underscoring the importance of BCAA metabolism for adipocyte metabolic flexibility^47^. Additionally, we found that isovaleryl-CoA dehydrogenase (IVD) mRNA levels are reduced in our dataset; notably, SNPs in the IVD gene are associated with T2D and regulate its expression. An earlier study demonstrated that leucine catabolism increases thirtyfold during 3T3-L1 adipogenesis, with IVD activity rising twentyfold. However, the role of IVD in adipocytes has received little attention since that time^48^. Collectively, our results reinforce the idea that BCAA metabolism is a key determinant of adipocyte homeostasis. Future research should aim to clarify how BCAA pathways influence adipose tissue function and, potentially, whole-body metabolism.

Beyond BCAAs, our data also reveal that genes involved in the metabolism of cysteine, proline, glutamine, and tryptophan are repressed in adipose tissue across both of our models. Among these, CdO1, which catalyzes the transformation of cysteine to taurine, stands out. Recent work has shown that CdO1 regulates lipolysis in adipocytes, and its deletion exacerbates the harmful effects of a high-fat diet in mice. Mechanistically, CDO11 appears to exert its effects not through taurine metabolism, but via physical interaction with PPARγ and regulation of ATGL and HSL activity^49^. Thus, while our data support that CDO1 plays an important role in adipocyte homeostasis, it remains to be determined if this is, at least partially, due to its function in cysteine metabolism. PRODH, a gene associated with proline oxidase activity, has also been linked to the regulation of lipolysis in adipocytes, although its precise metabolic function remains to be defined^50^. Finally, the alcohol dehydrogenase, iron-containing protein 1 (ADHFE1), has been little studied in adipose tissue. Its mRNA levels increase during 3T3-L1 differentiation^51^, and interestingly, ADHFE1 is overexpressed in breast cancer, where it promotes tumor cell survival by enhancing metabolic flexibility and supporting reductive glutamine metabolism for citrate synthesis^52^. Given that ADHFE1 confers metabolic flexibility in cancer cells, one could speculate that its decreased expression in adipocytes might contribute to the metabolic inflexibility observed in obesity. This intriguing possibility warrants further functional investigation. Interest in ADHFE1 is further strengthened by our finding that SNPs in ADHFE1 are associated with T2D, and that eQTL analysis indicates these SNPs regulate its expression levels. In summary, our findings highlight the central role of amino acid metabolism in adipocyte function and insulin sensitivity, and point to several promising avenues for future research into how these pathways govern adipose tissue and systemic metabolic health.

Altogether, our work identifies a genuine molecular signature of adipocyte dysfunction. We show that dysfunctional adipocytes lose their capacity to utilize glucose and amino acids and to synthesize lipids from their catabolic products. Our approach has highlighted pathways and targets previously studied in the field, and the fact that our unbiased analysis converges on these genes and pathways reinforces their fundamental importance in maintaining adipocyte health. Nonetheless, a major challenge remains: integrating these findings into a holistic understanding of adipocyte dysfunction. How the collective downregulation of the genes identified here leads to impaired adipocyte function, and how this dysfunction contributes to the development of metabolic complications such as type 2 diabetes—seen in both obesity and lipodystrophy—remains to be fully elucidated. We believe, however, that by comparing two extremes of adipose tissue abnormalities, our study has narrowed the focus to 102 genes that may hold the key to unraveling the mechanisms underlying adipose tissue failure.

In conclusion, our findings demonstrate that a shared “energetic collapse” of adipocytes, characterized by profound metabolic inflexibility in pathways spanning glucose utilization, lipogenesis, and amino acid catabolism, represents a common pathogenic mechanism underlying adipose tissue dysfunction in both obesity and lipodystrophy. This convergent molecular signature underscores the critical role of intrinsic adipocyte metabolic health in systemic energy homeostasis and insulin sensitivity. By unveiling these shared dysregulations and identifying novel, genetically supported targets, our work opens new avenues for therapeutic interventions aimed at restoring adipocyte metabolic fitness to combat cardiometabolic diseases. Future functional studies are essential to elucidate the precise causal contributions of these identified pathways and genes to systemic insulin resistance, paving the way for targeted strategies to prevent and treat metabolic dysfunction associated with impaired adipose tissue.

## Materials and Methods

### Animal models

*Bscl2*lox/lox was established at the Institut Clinique de la Souris (http://www.phenomin.fr). Three clones were ordered from the European Mouse Mutant Cell Repository (https://www.eummcr.org/). All three clones were validated by Southern blot (internal Neo probe) and PCR confirmation of the presence of the 3’ LoxP site. Clone HEPD0757_2_A10 was karyotyped by chromosome spreading and Giemsa staining and microinjected in BALB/cN blastocysts. Resulting male chimeras were obtained and germ line transmission was achieved for the tm1a allele (knock-out first with conditional potential). The KO allele (tm1b) was obtained after breeding tm1a animals with a Cre deleter line (Birling et al., 2012) and phenotyped by PHENOMIN-ICS. All data are freely available through the IMPC web site (https://www.mousephenotype.org/data/genes/MGI:1298392). The cKO Bscl2 allele (tm1c; Lox/Lox) was obtained by breeding the tm1a allele with a Flp deleter line (Birling et al., 2012). Bscl2lox/lox mice were crossed with ERT2-Adipoq-CRE (Sassmann et al., 2010) mice to generate Bscl2 lox/lox mice X ERT2-Adipoq-CRE mice. Mice were housed at 21°C with a 12:12 h light-dark cycle with free access to food and water. CRE activation was performed by 5 days intraperitoneal injection of tamoxifen resuspended in sunflower oil: ethanol (9:1) to 8-week-old males. For the 3-month group, one intraperitoneal injection was repeated every month. Glucose tolerance test and Insulin tolerance test were performed after 6 hours fasting as previously described (Prieur et al., 2013). All mice (males, n = 8 to 11 per group) were euthanized in the random fed state between 8 a.m to 10 a.m. The ethics committee of the French national veterinary agency approved all animal protocols used in this study. Food was removed 6 h before the initiation of the intraperitoneal glucose tolerance test (IPGTT) or the insulin tolerance test (ITT). At time 0, a single dose of glucose (2 g/kg) or insulin (0.75 U/kg, Actrapid, Novo Nordisk) was injected i.p. and blood glucose levels were monitored using a glucometer (Freestyle Papillon, Abbott, France) on 2.5 ml samples collected from the tail.

For obesity model, male C57BL/6J mice were fed a 60% fat diet (HFD) for 1, 3, or 6 months.

### Methods details

3′-seq RNA profiling was performed as described previously (Cacchiarelli et al., 2015). Libraries were prepared from 10 ng of total RNA in 4 μL. mRNA poly(A) tails were tagged with universal adapters, well-specific barcodes, and unique molecular identifiers (UMIs) during template-switching reverse transcription. Barcoded cDNAs from multiple samples were pooled, amplified, and fragmented using a transposon-based approach (Nextera DNA Sample Prep Kit, Illumina, ref. FC-121-1030) that enriches for 3′ ends of cDNA. A total of 100 ng of full-length cDNA was used as input.

Library size was assessed using the 2200 TapeStation System (Agilent Technologies). Libraries ranging from 350 to 800 bp in length were sequenced on an Illumina HiSeq 2500 using the HiSeq Rapid SBS Kit v2 (50 cycles, ref. FC-402-4022) and the HiSeq Rapid PE Cluster Kit v2 (ref. PE-402-4002), according to the manufacturer’s protocol (*Denaturing and Diluting Libraries for the HiSeq and GAIIx*, Part #15050107 v03, Illumina).

Raw FASTQ read pairs met the following criteria: the first 16 bases of Read 1 consisted of a 6-base sample-specific barcode followed by a 10-base unique molecular identifier (UMI). Read 2 (58 bases) corresponded to the captured poly(A)+ RNA sequence.

We performed demultiplexing of the read pairs to generate one single-end FASTQ file per sample (n = 96). These FASTQ files were aligned using BWA to reference mRNA RefSeq sequences and to the mitochondrial genome, both obtained from the UCSC download site.

### Statistical analysis

Analysis of 3’seq-RNA profiling

Gene expression profiles were generated by parsing alignment files (.bam) and quantifying unique molecular identifiers (UMIs) associated with each gene. Reads mapping to multiple genes, containing more than three mismatches to the reference sequence, or displaying poly(A) patterns were discarded. This process yielded a matrix representing the absolute mRNA abundance of all genes across all samples, suitable for downstream gene expression analysis.

### Transcriptomic analysis

Differential expression analysis was performed using the DESeq2 R package (Love et al., 2014). To identify genes with significant time-dependent expression changes between experimental groups, we applied MaSigPro ^53^ to the log-transformed expression matrix. Using default parameters and a quadratic regression model, we first identified genes with expression profiles significantly altered compared to the CRE. Tamox control group in the lipodystrophy model and to the HFD group in the obesity model. To control for false positives, we applied the Benjamini–Hochberg correction (Benjamini and Hochberg, 1995), setting the false discovery rate (FDR) threshold at 0.05. A backward stepwise regression procedure was then used to refine the best-fitting model for each gene, yielding 653 differentially expressed genes in the lipodystrophy model and 1,007 in the obesity model. These genes were subsequently grouped into clusters based on similarities in their temporal expression profiles.

Public datasets included GSE97271 (microarray), GSE213632, E-MTAB-8840, and GSE167264 (RNA sequencing). GSE97271 represents the micro-array analysis of the gonadal AT from 6 C57BL6J male mice fed a HFD or a chow diet for 20 weeks. GSE213632, E-MTAB-8840 and GSE167264 are transcriptomic profiles obtained by RNA sequencing after, respectively, after 11, 16 and 20 weeks of HFD. Differential gene expression in these datasets was identified using the R packages LIMA ^54^(for microarrays) and edgeR ^55^ (for RNA-Seq).

MEGENA (Multiscale Embedded Gene Co-expression Network Analysis) tool (PMID: **26618778)** was utilized to construct gene-gene interaction networks and identify co-expression modules. To assess the correlation between gene expression and insulin resistance, Weighted Gene Co-expression Network Analysis (WGCNA) was performed (PMID: **19114008)**. Glucose tolerance test (GTT AUC) and insulin tolerance test (ITT AUC) values were used as phenotypic traits. Gene modules with expression significantly correlated, either positively or negatively, with AUC values were identified.

### Functional Enrichment Analysis

To explore the biological relevance of the overlapping gene signature, Gene Ontology (GO) enrichment analysis was conducted using ToppFun (from the ToppGene Suite). Analyses focused on the Biological Process and Cellular Component GO categories. Significantly enriched terms were identified using a hypergeometric test with Benjamini–Hochberg correction (FDR < 0.05), providing insight into the functional roles and subcellular localization of the deregulated gene sets.

### Literature Review and Gene Annotation

To contextualize the identified genes and assess their known relevance in the studied pathologies, a comprehensive literature review was performed. For each gene within our refined signature, we systematically searched the PubMed database (https://pubmed.ncbi.nlm.nih.gov/). Our search strategy involved combining the gene name with relevant keywords such as “obesity,” “adipose tissue,” “lipodystrophy,” and “adipocytes.” This approach allowed us to identify existing research and biological functions previously described for each gene in the context of adipose tissue dysfunction and related metabolic disorders.

### Correlation Analysis with Human Metabolic Traits

The relevance of our shared gene signature in humans was assessed by correlating the expression levels of our 102-gene list with key metabolic traits, including Body Mass Index (BMI), waist-to-hip ratio, HOMA-IR, HbA1c, glycemia, and insulin levels. For this analysis, we utilized the Adipose Tissue Portal (https://www.adiposeportal.org/), a publicly available resource compiling extensive adipose tissue transcriptomic data (PMID: **39983713**). Transcriptomic data specifically from human subcutaneous adipose tissue were used for this correlation analysis.

### Genome-Wide Association Analysis (GWAS) and eQTL

The genetic association with type 2 diabetes was assessed using summary statistics data from the cross-ancestry meta-analysis published by Suzuki et al. ^16^. The summary statistic files were downloaded via the DIAGRAM consortium portal (link 1). This study includes 428,452 type 2 diabetes (T2D) cases and 2,107,149 controls. The study population comprises six ‘ancestry groups’: European (60.3%); East Asian (19.8%); admixed African American (10.5%); admixed Hispanic (5.9%); South Asian (3.3%); and South African (0.2%).

Associations between genetic variants and mRNA expression levels (also known as expression quantitative trait loci (eQTL)) in adipose tissue were tested using a publicly available eQTL dataset called AdipoExpress^18^. This dataset comprises 2,344 subcutaneous adipose tissue samples. Link1: http://www.diagram-consortium.org Link 2: https://zenodo.org/records/13845120

---

**Supplemental figure 1.**
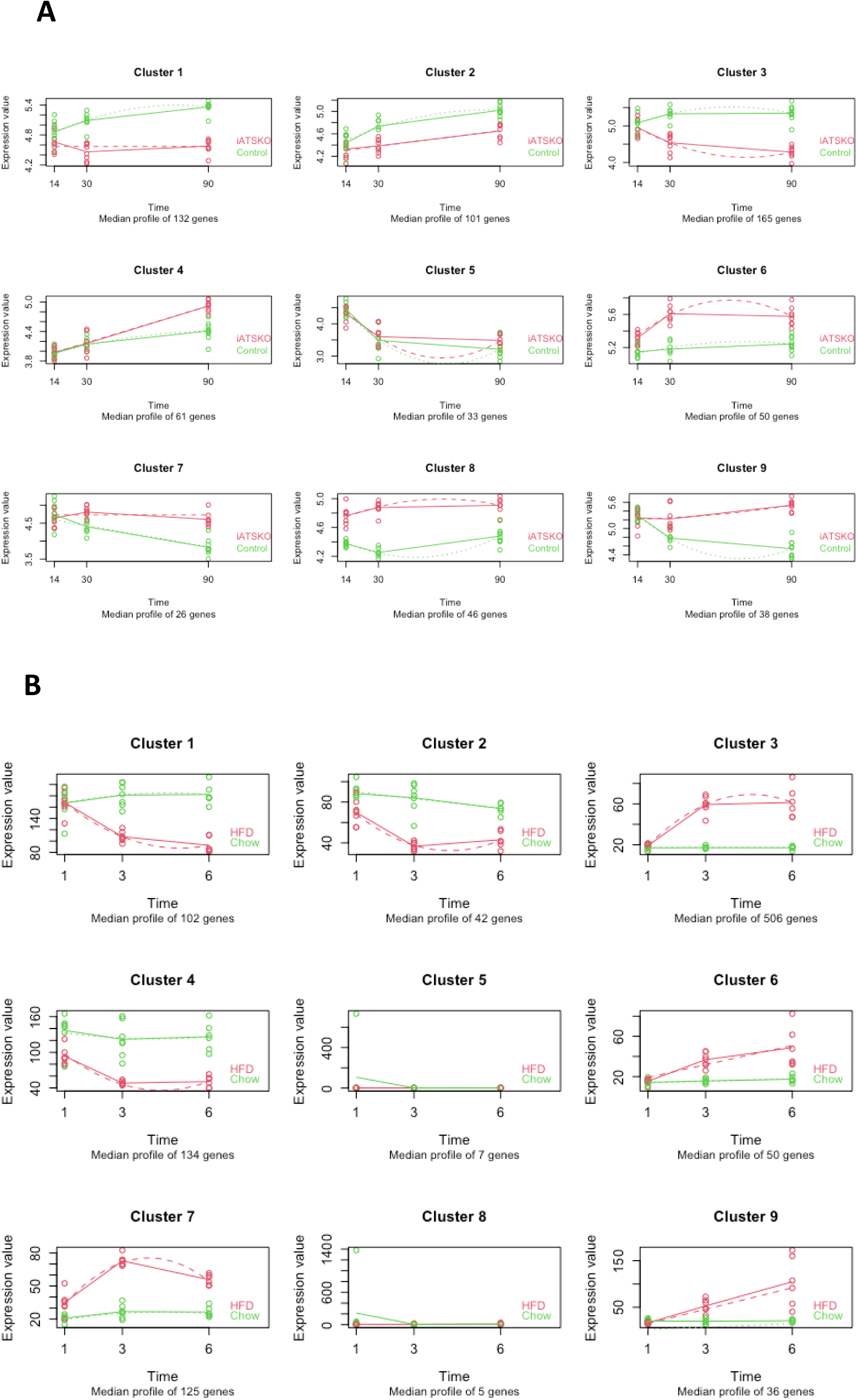
(A) Identification of Differentially Expressed Genes in Adipose Tissue of Lipodystrophic and Obese Mice. RNA sequencing analysis of subcutaneous adipose tissue from lipodystrophic iATSKO mice at 15 days (early), 30 days (middle), and 90 days (late) after tamoxifen injection. Gene expression data were clustered using MaSigPro, and the median expression profiles of the 9 clusters are shown over time. (B) RNA sequencing analysis of visceral adipose tissue from C57Bl/6J mice fed either a chow diet or a high-fat diet (HFD) starting at 8 weeks of age, with samples collected at 1 month (early), 3 months (middle), and 6 months (late). Gene expression data were clustered using MaSigPro, and the median expression profiles of the 9 clusters are shown over time.

**Supplemental figure 2.**
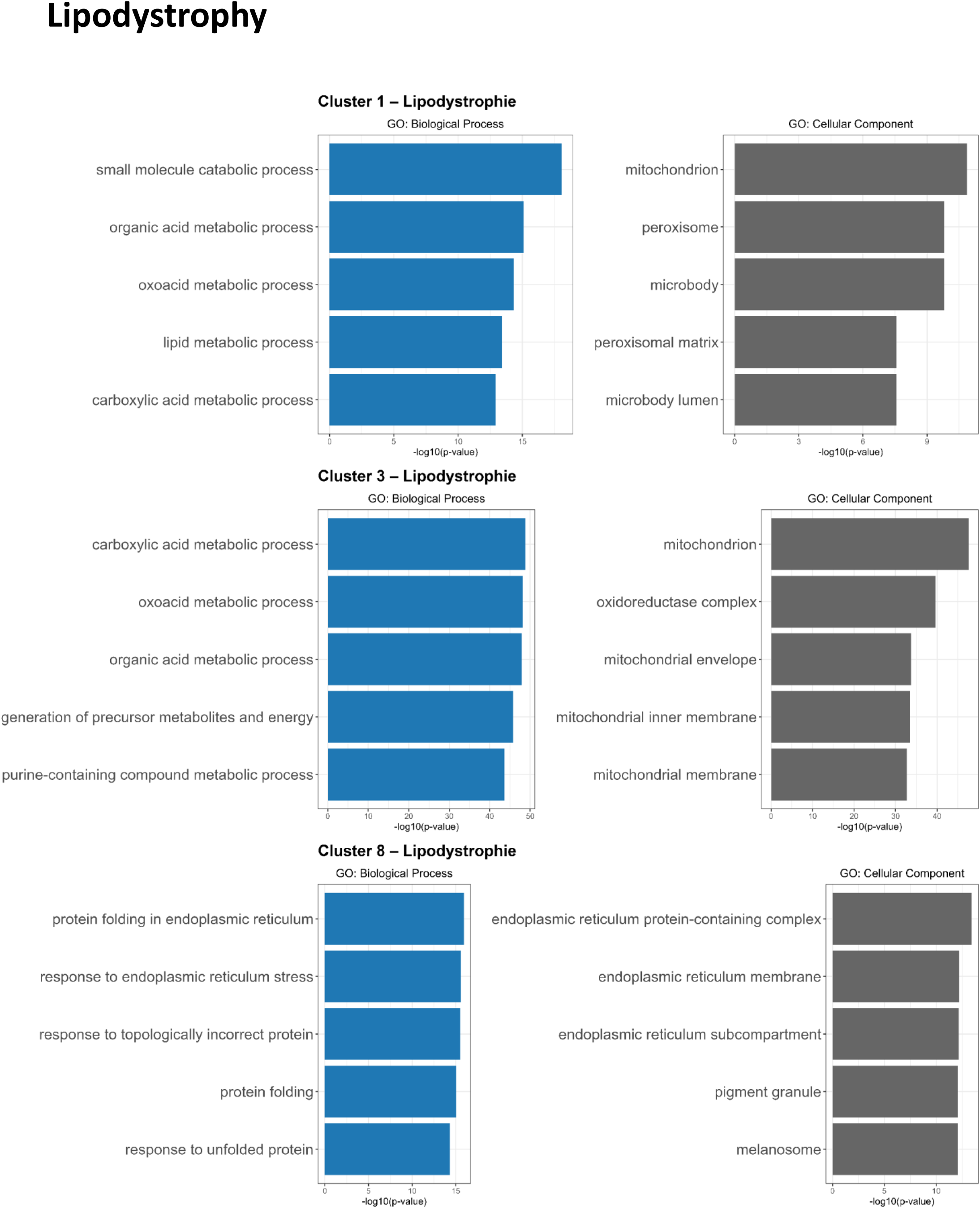
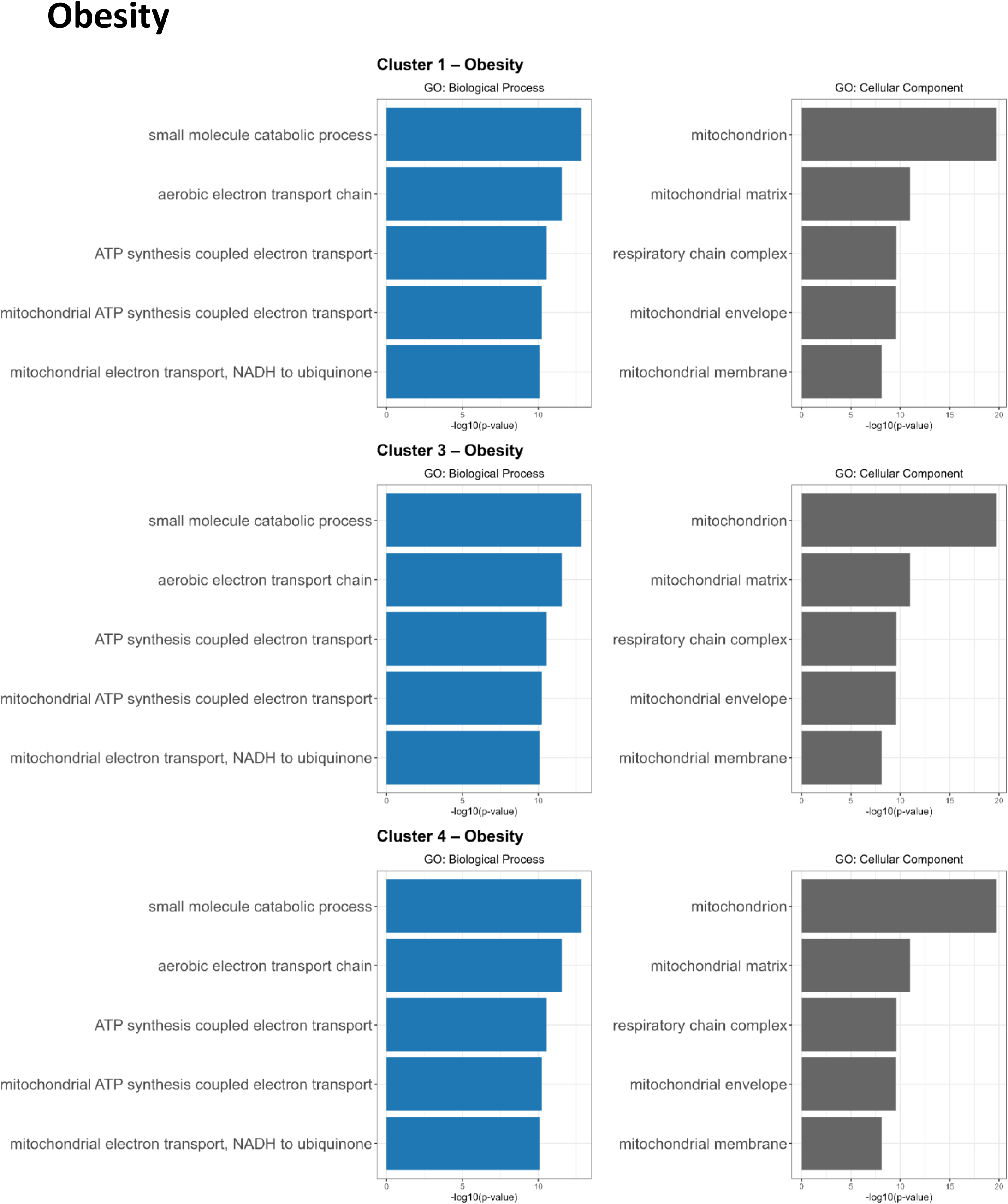
Gene ontology analysis of gene clusters identified by MaSigPro, starting from genes differentially expressed in both the lipodystrophy and obesity datasets. The table displays the main functional categories and cellular components enriched within these gene clusters.

**Supplemental figure 3.**
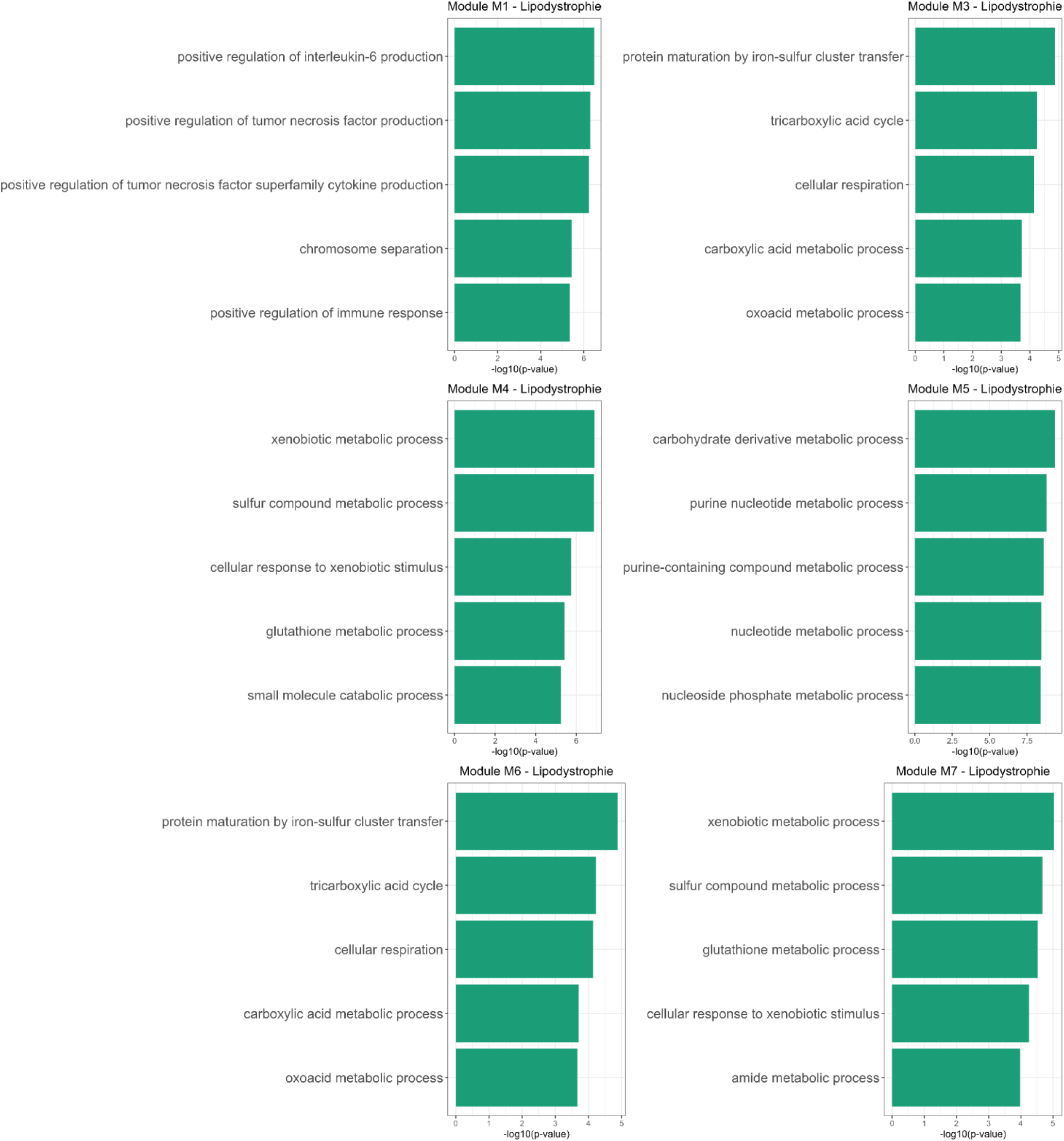
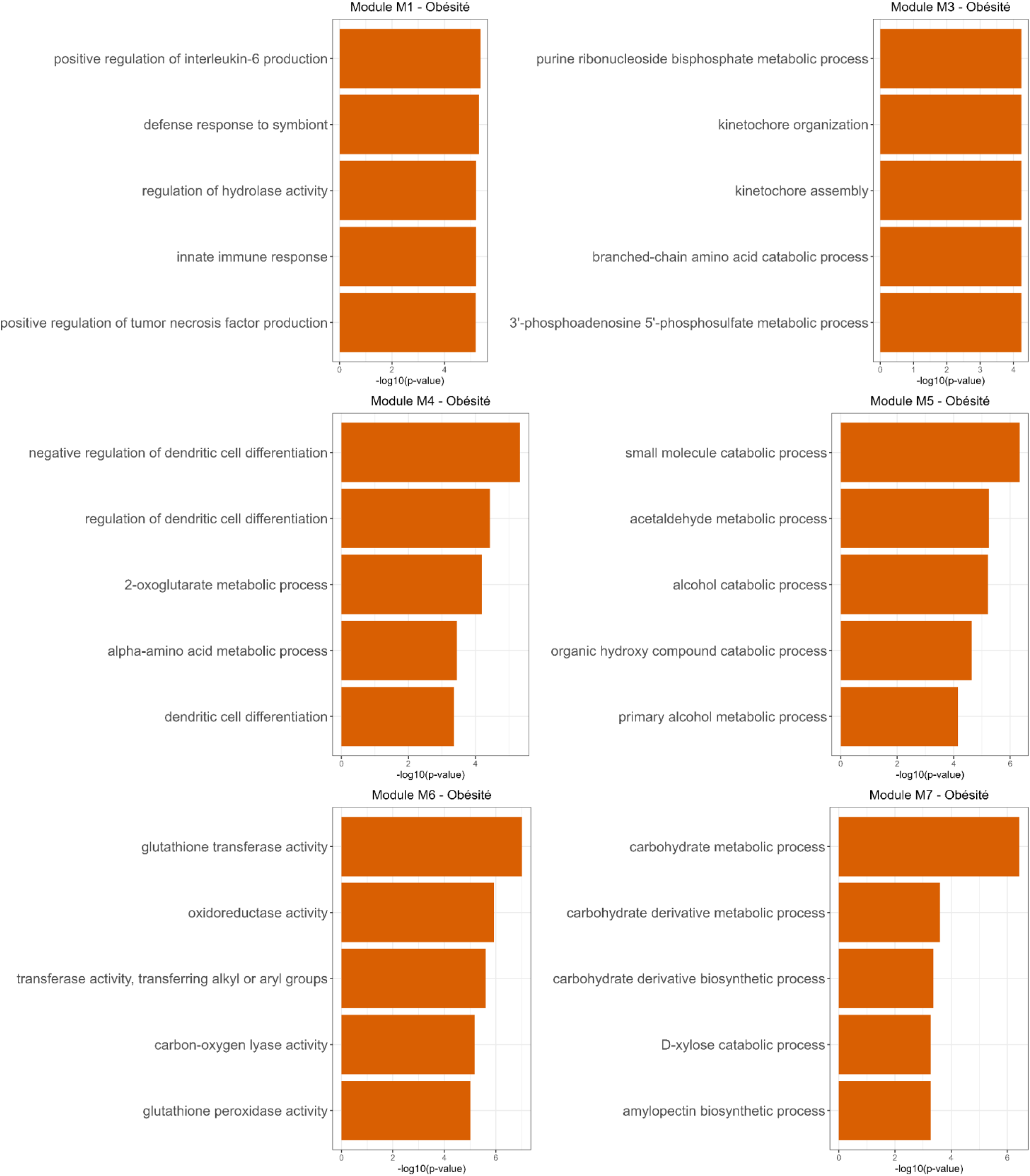
MEGENA (Multiscale Embedded Gene Co-expression Network Analysis) was used to construct gene-gene interaction networks and identify co-expression modules among the 129 genes in both the lipodystrophy and obesity datasets. Gene ontology analysis was performed in parallel on the modules generated from each dataset to explore enriched biological functions and pathways.

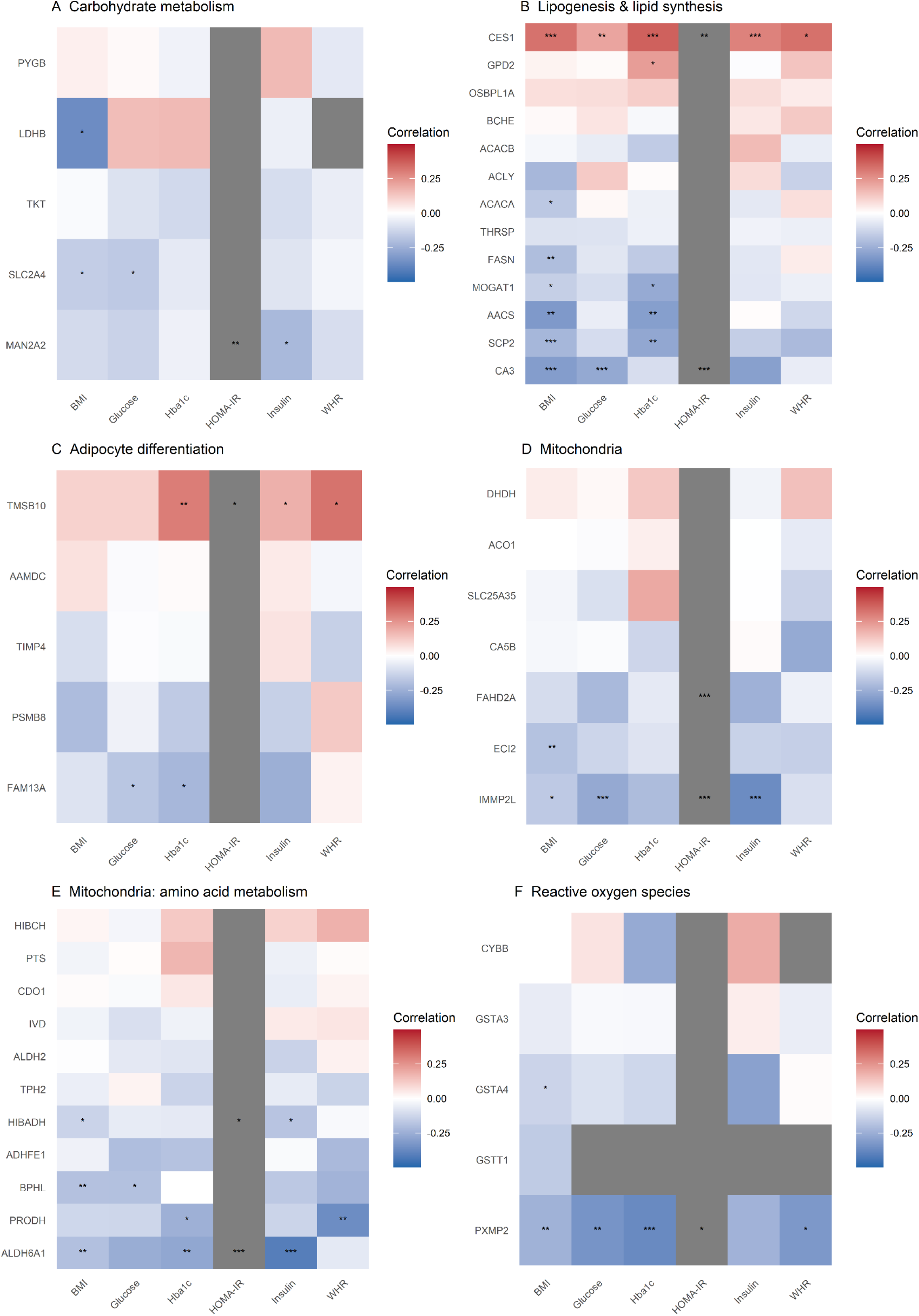
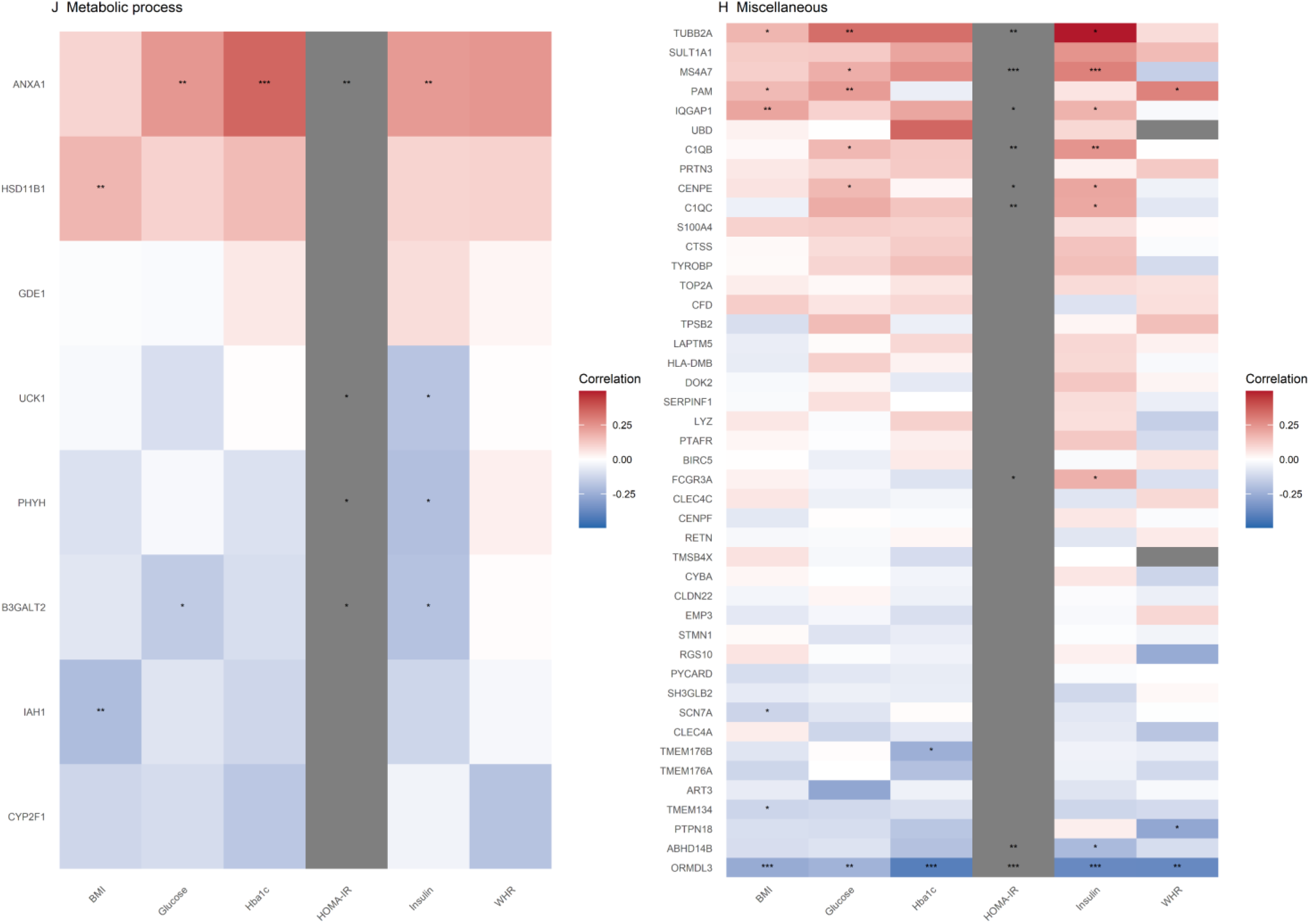

**Supplemental Table 1.**
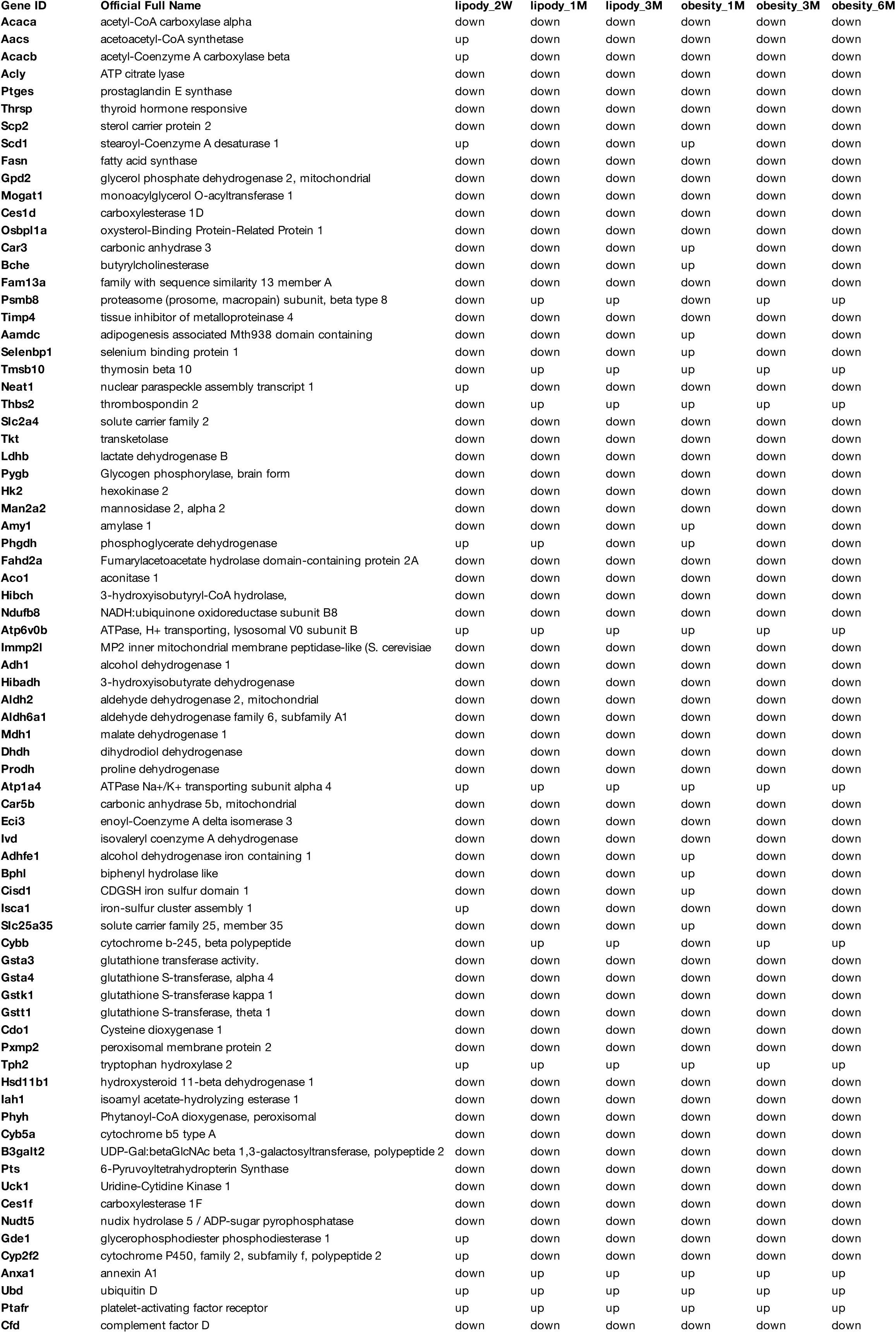
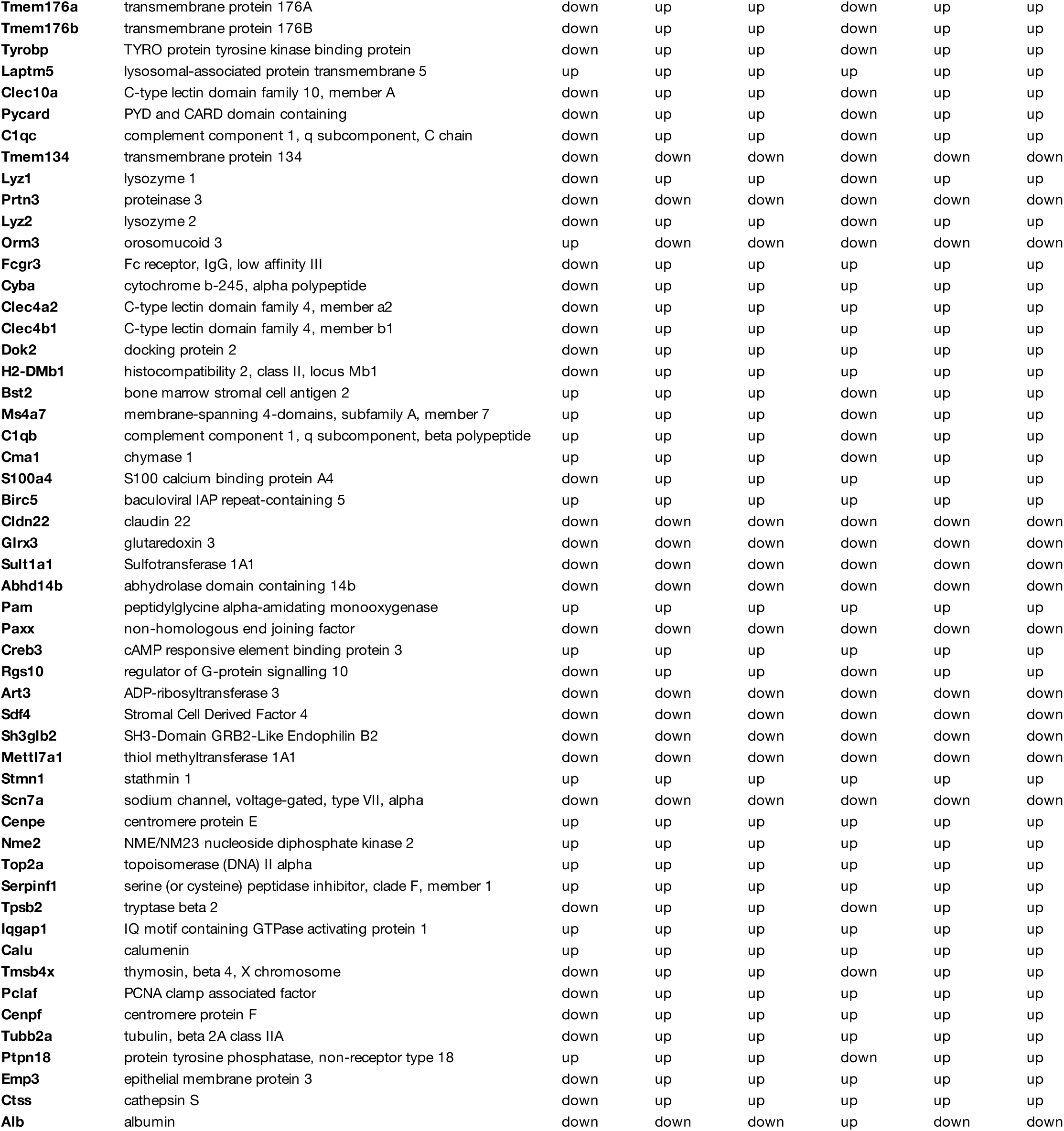

